# Enteral immunization with live bacteria reprograms innate immune cells and protects neonatal foals from pneumonia

**DOI:** 10.1101/2024.02.17.579919

**Authors:** Bibiana Petri da Silveira, Susanne K. Kahn, Rebecca M. Legere, Jocelyne M. Bray, Hannah M. Cole-Pfeiffer, Michael C. Golding, Noah D. Cohen, Angela I. Bordin

## Abstract

Neonatal immunization faces several challenges due to the immune system’s immaturity and naivety, and several vaccines that induce adequate immune response in adults fail to protect neonates. Using a horse foal model, we show that enteral administration of *Rhodococcus equi* to newborn foals overcomes neonatal vaccination challenges by reprogramming innate immune responses, inducing *R. equi*-specific adaptive humoral and cell-mediated immune responses and protecting foals against experimental pneumonia challenge. Immunized foals exhibit changes in the epigenetic profile of blood monocytes, >1,000 differentially-expressed genes in neutrophils, and higher levels of *R. equi*-specific IgG_1_, IgG_4/7_, and T helper type 1 response. Together, our data indicate that early life exposure to *R. equi* in the gastrointestinal tract can induce enhanced innate immune responses, which may facilitate the development of adaptive immune responses and protect neonates against pneumonia through a previously unappreciated gut-lung axis.

**Teaser:** Enteral immunization of newborn foals with live bacteria enhances systemic innate immune responses through the gut-lung axis.

## Introduction

Vaccine development for neonates is challenging due to the intrinsic peculiarities of the immature innate and adaptive immune systems, such as bias towards anti- rather than pro-inflammatory responses to innate signals and limited T helper type 1 (Th1) differentiation (*1*). Neonates from many species, including foals, manifest a functional deficiency of innate immune cells, including ineffective antigen presentation, primarily due to reduced expression of toll-like receptors, antigen-presenting molecules, and costimulatory factors (*2–6*). Because antigen-presenting cells instruct lymphocytes to induce appropriate immune responses (*7*), these deficiencies of the innate immune system also compromise the development of humoral and cell-mediated responses by neonates. Immunization strategies using administration of live attenuated vaccines by the mucosal or oral route appear to be superior in overcoming the challenge of achieving sufficient immunogenicity to develop innate and adaptive immunity in newborns (*8*). Three vaccines are licensed for use in human newborns: bacillus Calmette-Guérin (BCG), hepatitis B vaccine, and oral poliovirus vaccine (*8*). Interestingly, 2 of these vaccines - BCG (*9*) and oral poliovirus (*10*) - induce non-specific protective effects against other infectious agents and enhanced innate immune response. Stimulation of enhanced innate immune responses soon after birth are likely critical for adequate maturation of the immune system and response to vaccines because this period is concomitant with exposure to environmental microbes that drive maturation of the immune system (*11*); however, this process remains poorly understood in neonates.

*Rhodococcus equi* is a facultative intracellular bacterium that causes severe pyogranulomatous pneumonia in horse foals (*12, 13*). This pathogen also affects immunocompromised people and shares many features with *Mycobacterium tuberculosis* infection (*12, 13*). The ability of *R. equi* to replicate within macrophages and cause disease in foals is attributed to a plasmid that contains a pathogenicity island encoding the virulence-associated protein A (VapA) (*14*). Foals are more susceptible to *R. equi* infection in the first weeks after birth (*15, 16*), possibly because of both immaturity and naivety of the innate and adaptive immune systems (*2, 3, 17–21*). Despite decades of research, several vaccines using a diverse range of technologies have failed to protect foals against *R. equi* pneumonia (*22–26*). The only active immunization strategy that has consistently protected foals against intrabronchial infection is the enteral administration of live virulent *R. equi* (*27–29*). In other respiratory infections, the gut microbiota can play a protective role by improving mucosal immune responses in the lung through a process named gut-lung axis, specifically involving innate immune responses (*30–32*). Previous studies using enteral immunization of foals have only focused on adaptive immune responses, such as *R. equi*-specific humoral and cell-mediated immune responses (*28, 33, 34*). The role of innate immune responses of neonatal foals to enteral immunization in protecting against pneumonia, however, has not been systematically investigated.

Like humans, foals exhibit wide genetic variation, enabling clinical researchers to examine a diverse range of patient immunological responses and disease severity in an outbred population (*35*). Consequently, the foal model represents a powerful tool for studying the role of innate immunity in immune responses to vaccines in neonates. Here, we demonstrate the involvement of enhanced innate immune responses in the protection induced by enteral immunization of foals, shedding light on how this approach overcomes the limitations of vaccinating newborns. We show that enteral immunization induced epigenetic reprogramming of monocytes and altered the transcriptional profile of neutrophils, including the regulation of genes associated with the regulation by interferon gamma (IFN-γ), antigen presentation, and enhanced innate immunity. Moreover, enteral immunization stimulated *R. equi*-specific IgG_1_, IgG_4/7_, and Th1 responses. Together, our findings indicate that enteral immunization of neonatal foals induced innate immune cell reprogramming that generates a protective response against pneumonia.

## Results

### Enteral immunization of newborn foals protects against *R. equi* intrabronchial challenge

We gavaged foals with either saline (control group; n=6) or 2 × 10^10^ CFU of *R. equi* (immunized group; n=5) at ages 1 and 3 days (Fig. 1A). No foals from either group showed adverse effects after gavage. At age 28 days, we infected all foals intrabronchially with 2 × 10^6^ CFU of *R. equi*. Enteral immunization significantly (P = 0.0454) reduced pneumonia incidence after *R. equi* intrabronchial challenge: all 5 immunized foals remained healthy whereas 67% (4/6) of control foals developed clinical pneumonia (Fig. 1B). Similarly, the incidence of pulmonary consolidations or abscesses detected by thoracic ultrasonography was significantly (P = 0.013) lower in immunized foals (Fig. 1C); none of the immunized foals exhibited sonographic thoracic lesions whereas we identified lesions in 83% (5/6) of control foals. We classified foals with pulmonary lesions and no clinical signs as having subclinical pneumonia, a frequent presentation in naturally infected foals (*12*).

**Fig. 1:**
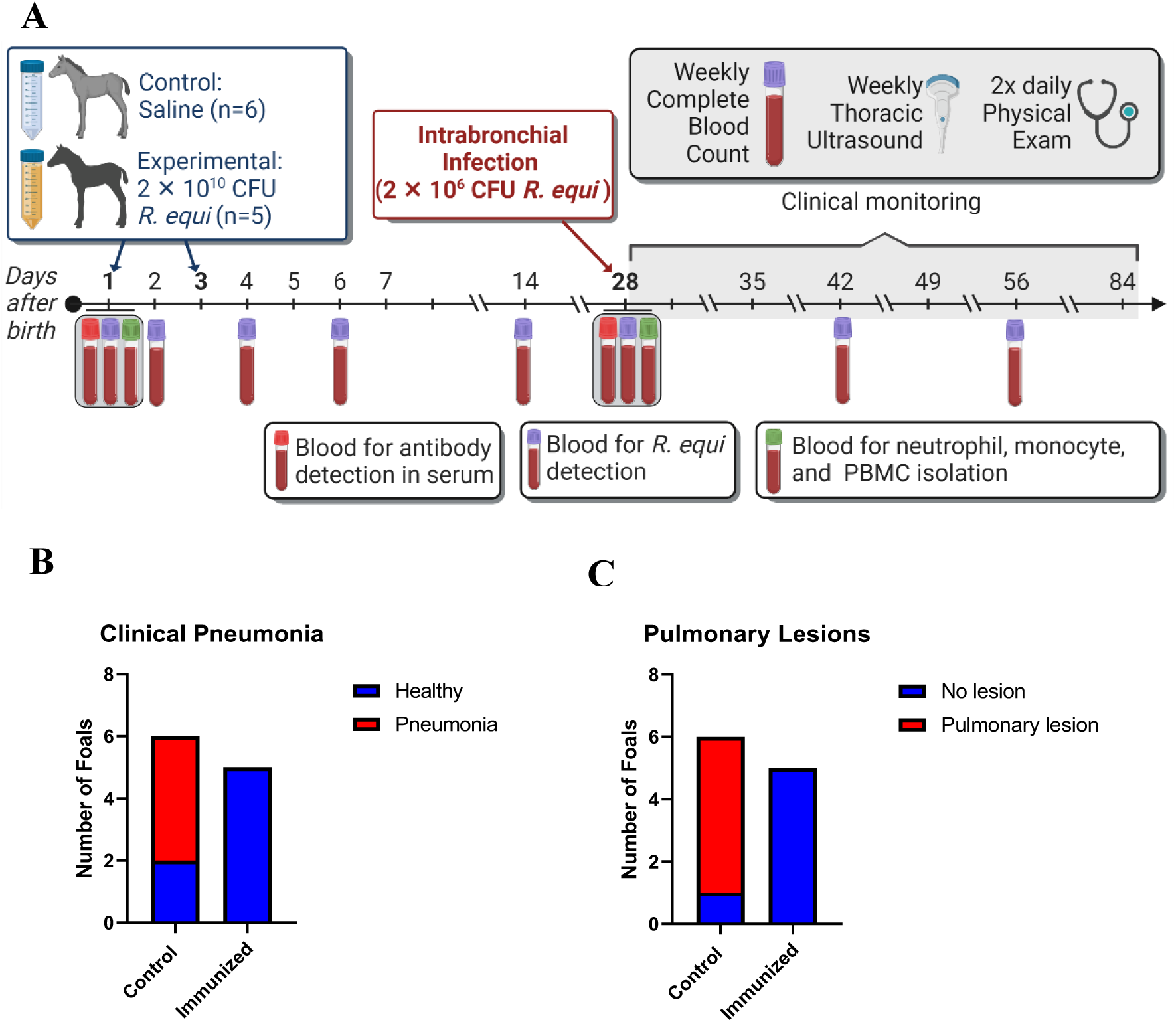
Enteral immunization of newborn foals protected against *R. equi* intrabronchial challenge at age 28 days. **A**) At ages 1 and 3 days, foals were gavaged with either 2 × 10^10^ CFU of *R. equi* (immunized group) or saline (control group). At age 28 days, all foals were challenged intrabronchially with 2 × 10^6^ CFU of *R. equi*. Blood for cell isolation was collected at ages 1 and 28 days. **B**) All 5 foals in the immunized group remained healthy after intrabronchial challenge with *R. equi*, in contrast to foals in the control group in which 67% (4/6) developed clinical pneumonia (P = 0.0454). **C**) Pulmonary lesions were detected by thoracic ultrasonography in 83% (5/6) in the control group, while no lesion developed in immunized foals (P = 0.013).

We could not isolate *R. equi* from the blood of immunized foals at all tested ages, suggesting that live bacteria do not reach the bloodstream following enteral immunization. In contrast, we isolated *R. equi* from blood of 50% (2/4) of the foals in the control group that developed pneumonia at age 42 days (2-weeks post-challenge, at the onset of clinical pneumonia). Both isolates were determined to be virulent based on positive results of PCR for the *vapA* gene. We believe *R. equi* isolated from sick foals was likely due to bacteremia following pneumonia caused by intrabronchial challenge.

### Enteral immunization with *R. equi* induces differentially-expressed genes in blood neutrophils

RNA-sequencing (RNA-seq) of blood neutrophils isolated from 1-day old foals before enteral immunization identified only 20 differentially-expressed genes (DEG; 1.5-fold change q<0.05) between the immunized and control groups; 17 were downregulated and 3 were upregulated (Fig. 2A). At 28 days, however, we found 1,152 DEG, with 248 downregulated and 904 upregulated in the immunized group compared to controls (Fig. 2A and 2B). The heatmap shows the genes with more than 11-fold-change between immunized and control groups, and the clustering of foals into groups of healthy foals (5 immunized and 1 control), and foals in the control group that developed clinical (n=4) or subclinical pneumonia (n=1) (Fig. 2B). These results suggest neutrophils from foals resistant to intrabronchial challenge had different gene expression profile compared to susceptible ones.

**Fig. 2:**
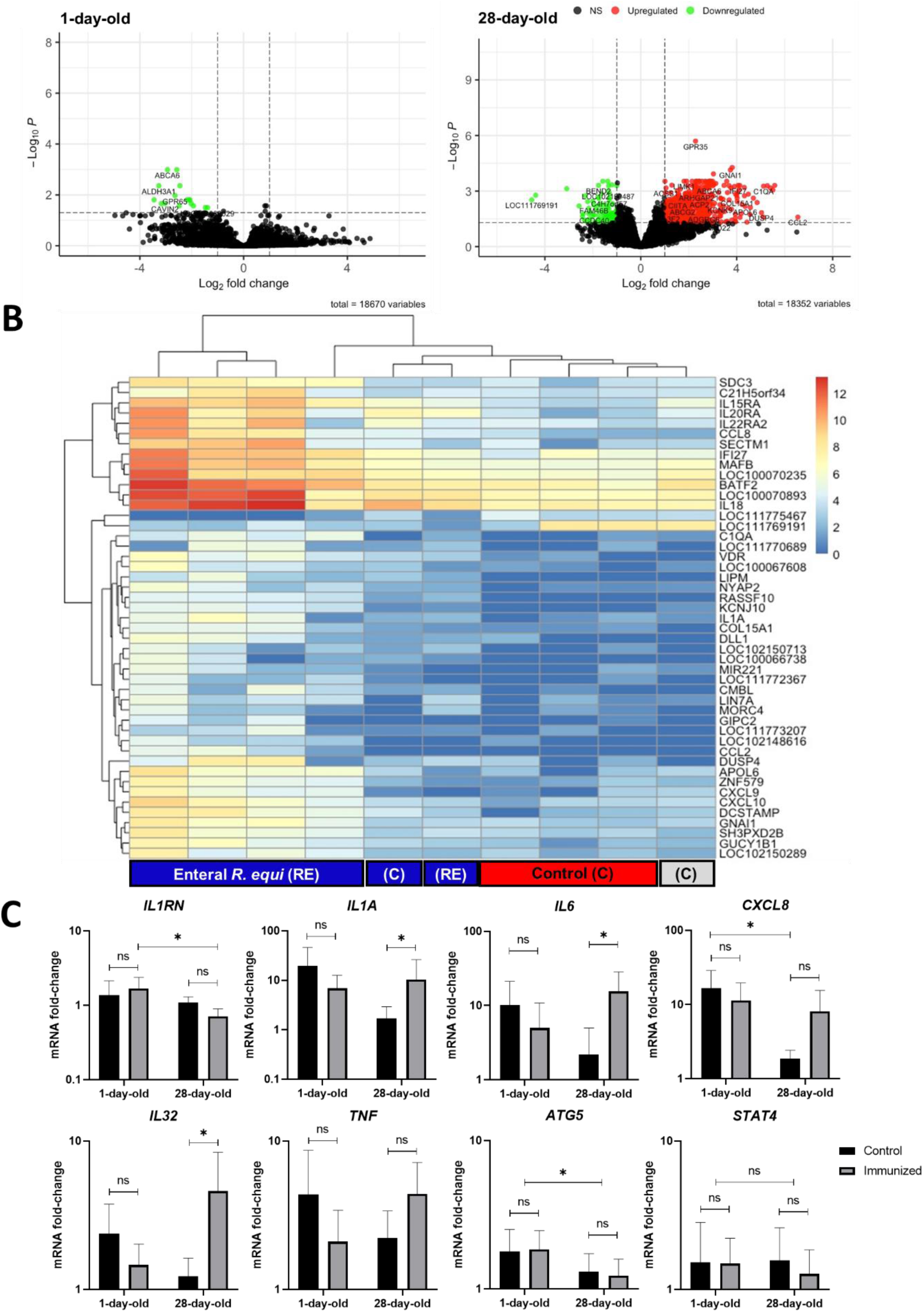
Enteral immunization with *R. equi* altered neutrophil gene expression. **A**) Volcano plot representation of RNA-Seq results showing differential expression analysis of genes in neutrophils of foals at ages 1 and 28 days between controls and immunized foals (dots represent difference on gene expression; ● = non-significant, ● = upregulated, ● = downregulated). **B**) Heatmap of RNA-Seq results of neutrophil from 28-day-old foals including all genes with fold-change higher than 11 between immunized and control groups (royal blue = healthy; red = pneumonic; gray = subclinical pneumonia). **C**) Gene expression results using qPCR. Fold-change of mRNA expression (y axis) at ages 1 (before immunization) and 28 (before intrabronchial challenge) days. Asterisk indicates significant statistical difference (p < 0.05) and ns (p > 0.05). Error bars represent standard deviation (SD).

Using qPCR, we confirmed neutrophils from immunized foals had higher expression of *IL1A*, *IL6*, and *IL32* at age 28 days (Fig. 2C). Expression of the *IL1RN* gene, which encodes an IL-1 receptor antagonist, decreased with age in the immunized group only, while *CXCL8* decreased in the control group only and remained high in the immunized group. There were no significant differences observed in the expression of *TNF*, *ATG5*, and *STAT4* genes between groups, but *ATG5* decreased significantly with age in both groups. Importantly, all those differences were detected in neutrophils collected from blood immediately before challenge, representing differences in their gene expression at basal levels before intratracheal infection.

Data from neutrophil gene expression indicated the immunized group at age 28 days had higher expression of pro-inflammatory cytokines (*IL1A*, *IL6*, *CXCL8*, *IL15*, *IL18*, *IL32*) and their receptors (*IL1R1*, *IL15RA*), trained immunity-related genes (*IL32*, *IL6*, *CD14*, *CXCL9*, *CXCL10*), proteins associated with antigen processing and presentation to T-cells (*CD1C*, *CIITA*, *CD74*, *PSMB9*, *PSMB10*, *PSME2*, *DRA*, *DRB*, *CD274*, *CTSA*, *CTSB*, *CTSC*, *CTSH*, *CTSZ, SECTM1*), regulation of the complement system (*C1QA*, *C1QB, LAIR1 MAFB*), hematopoiesis regulation (*CD74*, *DLL1, DCSTAMP, IL7, CEBPA, SECTM1*), leukocyte activation and response to microbes (*CD14, GNAI1, DUSP4, CCL2, CYBB, VDR*), cathepsins (*CTSA*, *CTSB*, *CTSC*, *CTSH*, *CTSZ*), and neutrophil clearance and homeostasis (*APOL6*, *CAPN2*, *C1QA*, *C1QB, BCL2A1*, *XAF1*, *NOD1*, *AIM2)*, among others.

Additionally, we found upregulation of several IFN-γ-induced genes such as *CXCL9*, *CXCL10*, *IL1A*, *CIITA*, *CD74*, *CD274*, *PSMB9*, *PSMB10*, *GBP5*, *NOD1*, *CYBB*, *APOL6*, *DRA*, *DRB*, *CTSA*, *CTSB*, *CTSC*, *CTSH*, *CTSZ*, *BATF2*, *BST2*, *CEBPA*, *NR4A1*, *APOL6*, *CAPN2*, *BCL2A1*, *XAF1*, *NOD1*, *AIM2, SECTM1, C1QA,* and *C1QB,* suggesting increased systemic IFN-γ stimulation (*36–39*). Using Ingenuity Pathway Analysis (IPA), we identified main pathways altered by enteral immunization of foals, including macrophage activation, mTOR signaling, HMGB1 signaling, cell death, cell movement, cell signaling, cell development, immune cells trafficking, hematological system development and function (Fig. 3). Importantly, IFN-γ was identified as an activated upstream regulator (P = 3.1E-25) (Fig. 3). For regulator effects and molecular networks, a consistency score of 3.33 indicated STING1 was the top regulator effect network affecting leukocyte movement and migration. Analysis of gene expression data revealed important changes on neutrophils induced by immunization, as well as genes and pathways potentially involved in protection against pneumonia.

**Fig. 3:**
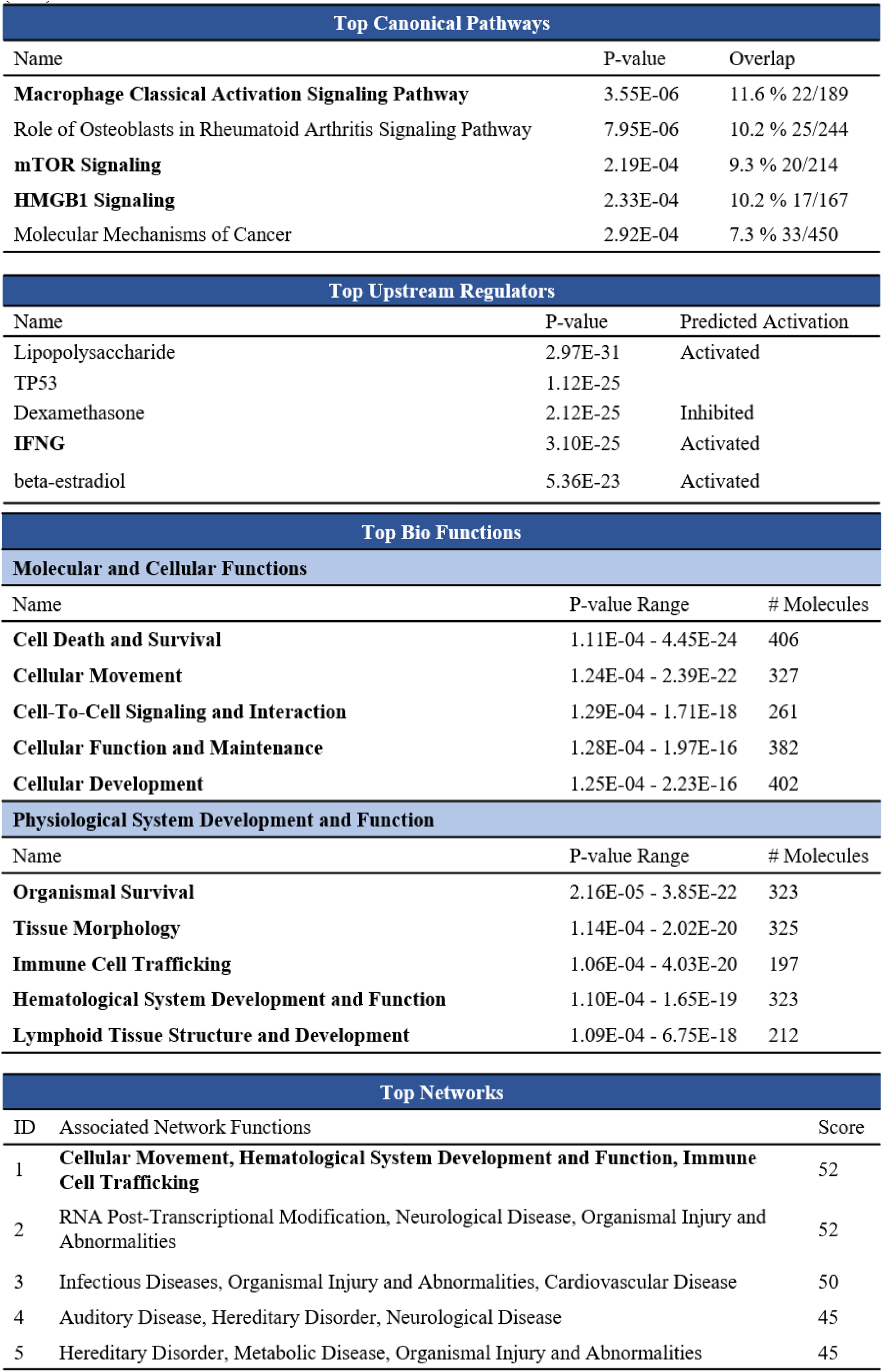
Top 5 pathways altered by enteral immunization in each category according to Ingenuity Pathway Analysis. Results of Ingenuity Pathway Analysis for differentially-expressed genes (DEG) in neutrophils from foals at age 28 days comparing immunized and control groups.

### Enteral immunization induces epigenetic reprogramming of circulating monocytes

From our transcriptomic data, we selected target genes with higher fold-change between groups at age 28 days, including *IL1A*, *C1QA*, *CCL8*, *CXCL8*, *CXCL10*, *SECTM1*, *VDR, IL1RN* and *IL32*, as well as other genes of biological interest such as *IL1B* and *TNF* (*40, 41*) for chromatin immunoprecipitation (ChIP) followed by qPCR (ChIP-qPCR). We performed ChIP assay on formaldehyde-fixed blood neutrophils and monocytes from foals at ages 1 and 28 days. Using monoclonal antibodies (mAb) and protein G magnetic beads, we isolated chromatin fragments associated with 2 histone modifications: the activating modification trimethylated histone 3 lysine 4 (H3K4me3) and the silencing modification histone 3 lysine 27 (H3K27me3). To perform qPCR, we purified DNA and designed primers specific for the region proximal to the transcriptional start site (TSS) of genes encoding *IL1A*, *IL1B*, *IL1RN*, *IL32*, *C1QA*, *CCL8*, *CXCL8*, *CXCL10*, *RPL30*, *SECTM1*, *TNF*, and *VDR*.

In monocytes, ChIP-qPCR analysis showed significant difference between groups (control vs immunized) for the % of input of H3K4me3. H3K4me3 increased significantly from ages 1 to 28 days in immunized foals for *IL1A*, *CXCL8*, *TNF*, and *RPL30* (P < 0.5; Fig. 4). In the control group, H3K4me3 decreased significantly for *VDR* (P < 0.05; Fig. 4). Comparing controls and immunized foals at age 28 days, immunized foals had higher H3K4me3 % of input for *C1QA, RPL30*, *SECTM1*, *VDR* (P < 0.05; Fig. 4). The percentage of input of target genes differentially enriched for H3K27me3 did not differ significantly between control and immunized groups. The only significant difference detected was a decrease in *IL32* with age, independent of group (Fig. S1). We also analyzed the ratio of H3K4me3:H3K27me3 because a higher ratio may indicate higher mRNA expression (*42*). In immunized foals, this ratio increased with age for *IL1A*, *IL1B*, *IL32*, *RPL30*, and *SECTM1* (P < 0.05).; however, in the control group we observed a decrease in the ratio for *C1QA* (P < 0.05) (Fig. 5) and no other significant changes. Comparing control and immunized foals at age 28 days, monocytes from immunized foals had a higher H3K4me3:H3K27me3 ratio for *SECTM1* (P< 0.05; Fig. 5).

**Fig. 4:**
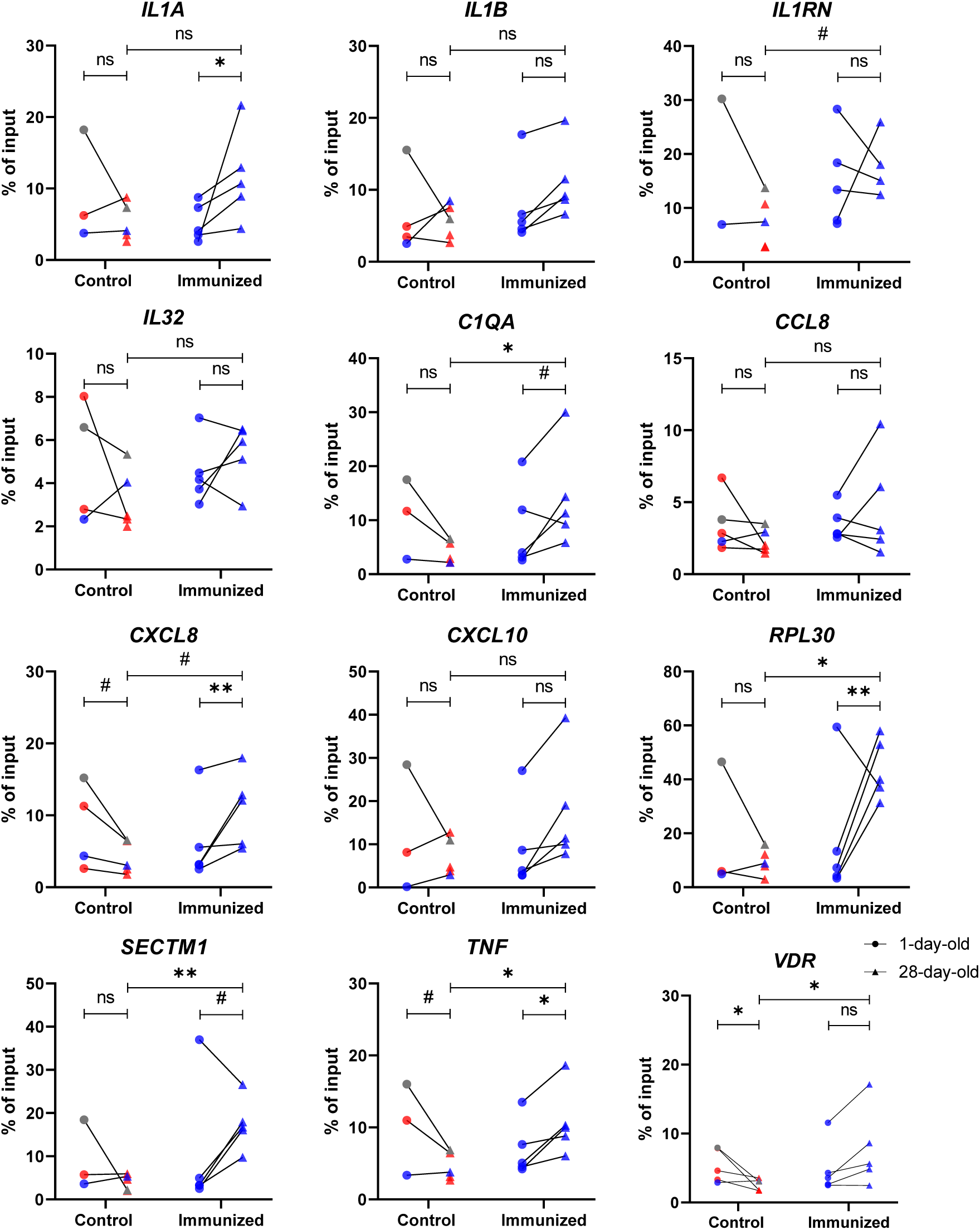
Enteral immunization induced H3K4me3 enrichment in the promoter regions of monocytes. Monocytes from immunized foals had an increase of H3K4me3 enrichment in the promoter regions at age 28 days. At ages 1 and 28 days, we isolated blood monocytes from foals and immunoprecipitated chromatin extracts with monoclonal antibody (mAb) recognizing the trimethylated form of histone 3 lysine 4 (H3K4me3). We purified DNA and performed qPCR using primers recognizing the region proximal to the transcriptional startsites of genes encoding *IL1A*, *IL1B*, *IL1RN*, *IL32*, *C1QA*, *CCL8*, *CXCL8*, *CXCL10*, *RPL30*, *SECTM1*, *TNF*, and *VDR*. Data are shown as % of input. Different symbols represent timepoints (● = 1-day-old and ▲ = 28-day-old). Different symbols represent different timepoints (● = 1-day-old and ▲ = 28-day-old). Blue symbols represent monocytes isolated from foals that remained healthy after intrabronchial infection, gray symbol the foal that developed subclinical pneumonia, and red symbols foals that had clinical pneumonia. The lines connect timepoints from the same animal. Statistical difference is represented by ** for P < 0.01, * for P < 0.05, # for 0.1 > P > 0.05, and ns for not significant.

**Fig. 5:**
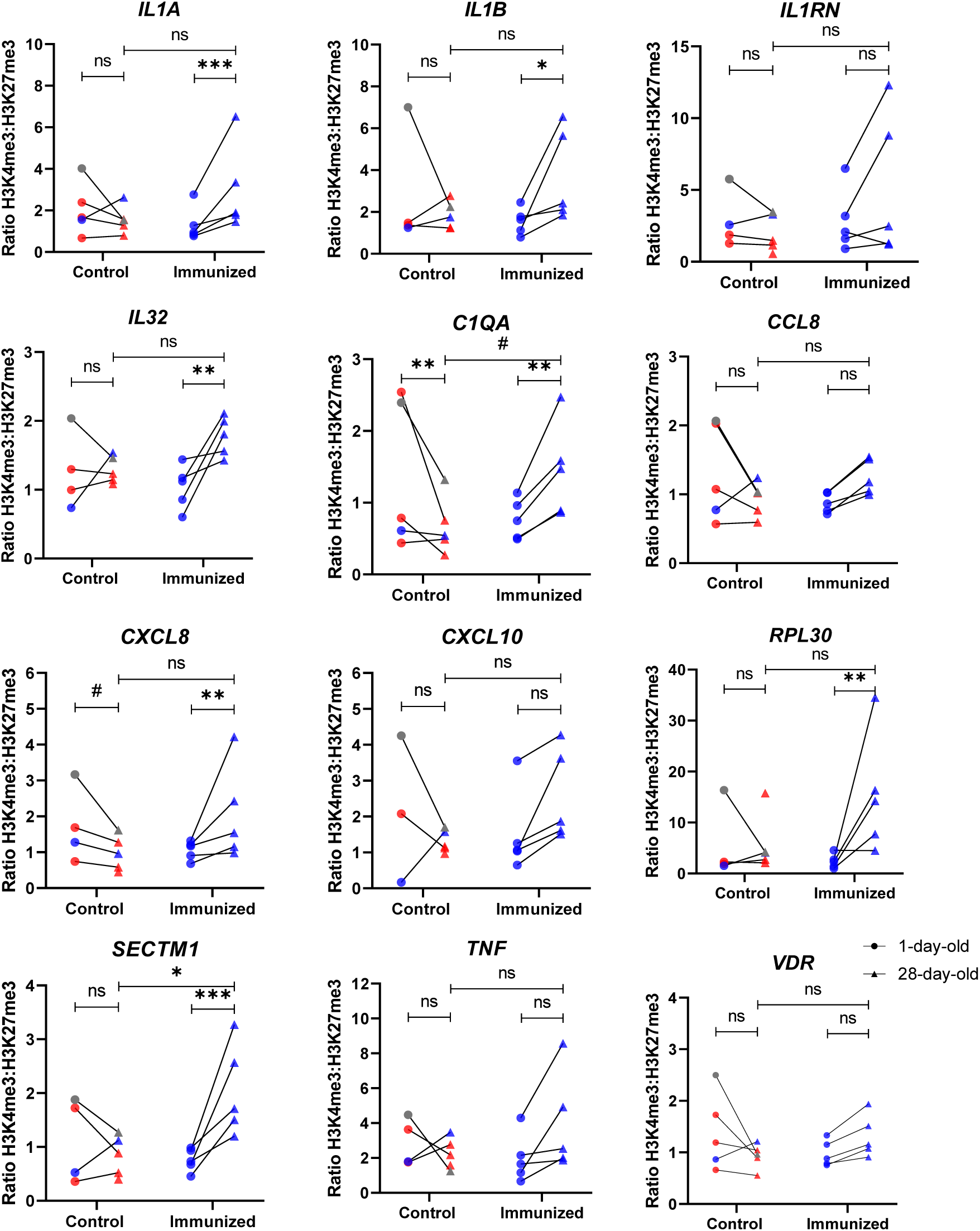
Enteral immunization increased H3K4me3:H3K27me3 ratio in the promoter regions of monocytes. We isolated blood monocytes from foals at ages 1 and 28 days. We immunoprecipitated chromatin extracts with mAbs recognizing the trimethylated forms of histone 3 lysine 4 (H3K4me3) and lysine 27 (H3K27me3). We performed qPCR using purified DNA and primers amplifying the regulatory regions of genes encoding *IL1A*, *IL1B*, *IL1RN*, *IL32*, *C1QA*, *CCL8*, *CXCL8*, *CXCL10*, *RPL30*, *SECTM1*, *TNF*, and *VDR*. Data show the ratio of % of input of H3K4me3 divided by % of input of H3K27me3. Different symbols represent timepoints (● = 1-day-old and▲ = 28-day-old). Different symbols represent different timepoints (● = 1-day-old and ▲ = 28-day-old). Blue symbols representmonocytes isolated from foals that remained healthy after intrabronchial infection, gray symbol the foal that developed subclinical pneumonia, and red symbols foals that had clinical pneumonia. The lines connect timepoints from the same animal. Statistical difference is represented by *** for P < 0.001, ** for P < 0.01, * for P < 0.05, # for 0.1 > P > 0.05, and ns for not significant.

In neutrophils, analysis of group, age, or group by age interaction did not reveal significant differences in enrichment of either H3K4me3 (Fig. S2) or H3K27me3 (Fig. S3) for the selected genes. Moreover, the H3K4me3:H3K27me3 ratio did not differ significantly between groups; however, the ratio increased with age in both groups for *IL1A*, *IL1B*, *CXCL8*, and *RPL30* (P < 0.05) (Fig. S4).

Interestingly, most of the genes with high fold-change in RNA-seq presented bivalent domains, characterized by enrichment of both H3K4me3 and H3K27me3 (Fig. 6). The genes for which we observed this bivalent domain in neutrophils were *IL32*, *C1QA*, *CCL8*, *SECTM1*, and *VDR*. Other genes such as *IL1B*, *IL1RN*, and *RPL30* had high H3K4me3 only. In monocytes, we also observed bivalent domains on the regulatory regions of *IL32*, *C1QA*, *CCL8*, *SECTM1*, and *VDR*; we also found bivalent domains for *IL1A*, *CXCL8*, and *CXCL10* (Fig. S5). Because of this commonality between monocytes and neutrophils, we believe the bivalent domain in the promoter regions of *IL32*, *C1QA*, *CCL8*, *SECTM1*, and *VDR* genes might be intrinsic to cells derived from myeloid progenitor cells in foals.

**Fig. 6:**
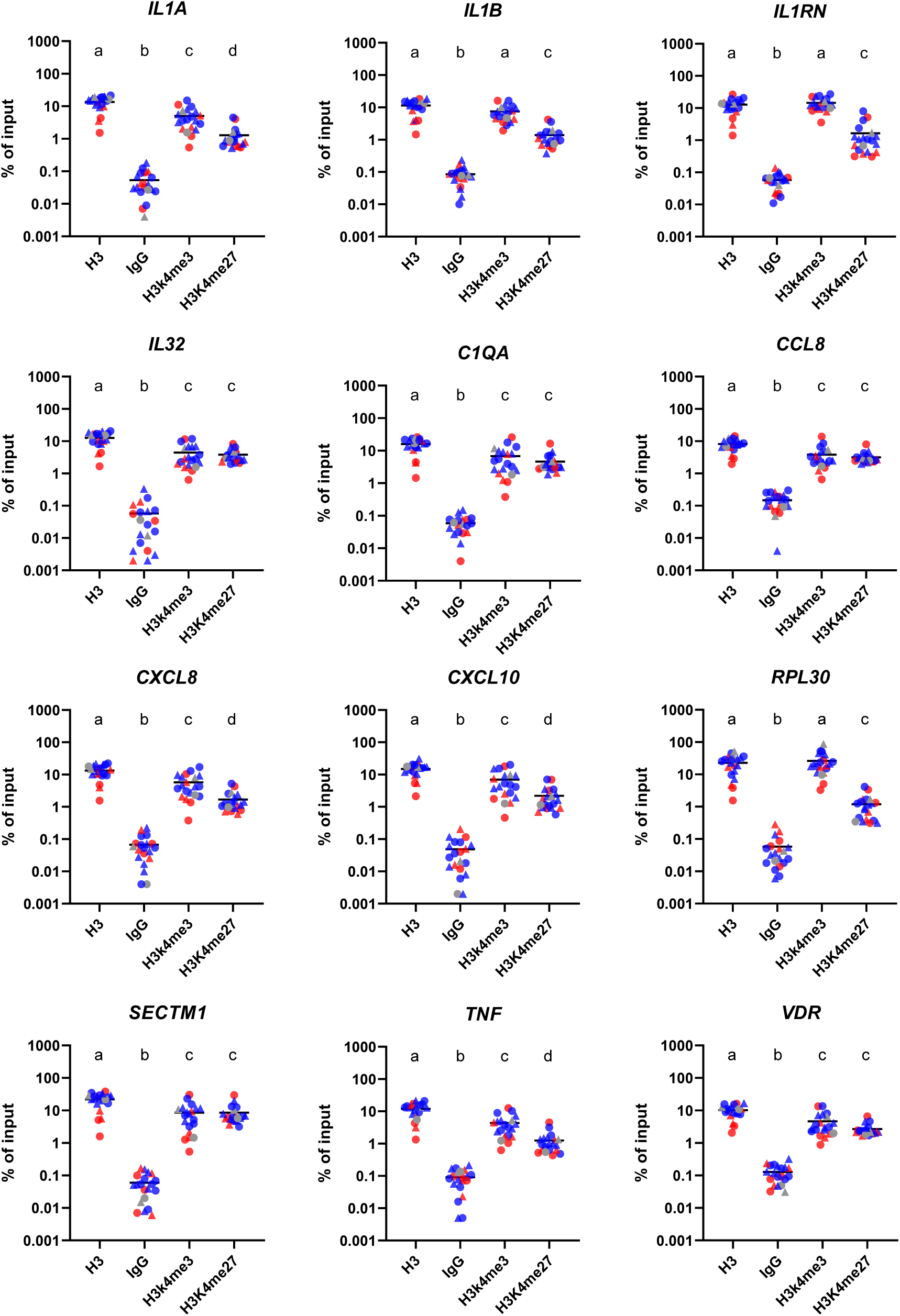
Neutrophils from foals exhibit a bivalent chromatin signature at genes encoding *IL32*, *C1QA*, *CCL8*, *SECTM1*, and *VDR.* We isolated blood neutrophils from all foals at ages 1 and 28 days. We immunoprecipitated chromatin extracts with mAbs recognizing the trimethylated forms of histone 3 lysine 4 (H3K4me3) and lysine 27 (H3K27me3), and the normal H3 histone (H3), normal IgG (mock control). We performed qPCR using primers specific for the region proximal to the transcriptional start site of genes encoding *IL1A*, *IL1B*, *IL1RN*, *IL32*, *C1QA*, *CCL8*, *CXCL8*, *CXCL10*, *RPL30*, *SECTM1*, *TNF*, and *VDR*. Bivalent domains were identified as those, having enrichment for both H3K4me3 and H3K27me3. Different symbols represent different timepoints (● = 1-day-old and ▲ = 28- day-old). Blue symbols represent neutrophils isolated from foals that remained healthy after intrabronchial infection, gray symbol the foal that developed subclinical pneumonia, and red symbols foals that had clinical pneumonia. Different letters (a, b, c, and d) represent statistical difference (P < 0.05).

### Enteral immunization resulted in *R. equi*-specific antibodies and Th1 immune response

At age 1 day, before enteral immunization or saline administration, foals had low levels of both VapA-specific IgG_1_ and IgG_4/7_, and we did not observe significant difference between groups (Fig. 7B,C). At age 28 days, activity of VapA-specific IgG of both subisotypes had increased in immunized foals while activities for control foals remained unchanged. To assess cell-mediated immunity, we isolated PBMCs from foals at age 28 days and stimulated them *ex vivo* with *R. equi* lysate to determine the number of spot-forming cells (SFC) that produced IFN-γ in an ELISpot assay. PBMCs from immunized foals had increased number of SFC than controls (p = 0.0168; Fig. 7A). In summary (Fig. 8), enteral immunization significantly enhanced *R. equi*-specific Th1 cell-mediated immunity (Fig. 7A), and increased VapA-specific antibodies in serum, both IgG_1_ (Fig. 7B) and IgG_4/7_ subtypes (Fig. 7C).

**Fig. 7:**
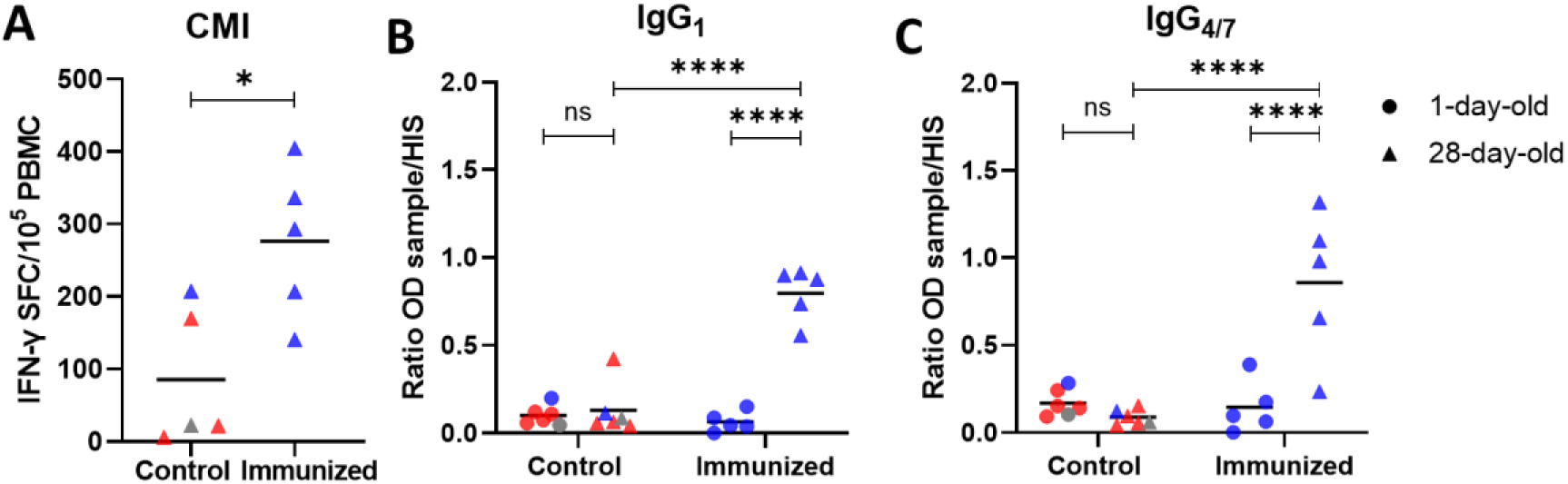
Enteral immunization induced *R. equi*-specific cell-mediated immunity (CMI) and humoral immunity. **A**) Peripheral blood mononuclear cells (PBMCs) isolated from foals at age 28 days that received enteral *R. equi* had increased (p = 0.0168) IFN-γ spot-forming cells (SFC) following stimulation with *R. equi* lysate, suggesting induction of higher *R. equi*-specific immune responses compared to control foals. **B** & **C**) Anti-VapA IgG_1_ (**B**) and IgG_4/7_ (**C**) activities were measured in foal serum at ages 1 and 28 days by ELISA. The ratio of the optical density (OD) for each sample was calculated by dividing the sample OD minus blank by the positive control *R. equi*-hyper immune serum (HIS) minus blank. Both VapA-specific IgG_1_ and IgG_4/7_ subisotypes were increased at age 28 days in immunized foals, but not in control foals. Different symbols represent timepoints (● = 1- day-old and ▲ = 28-day-old). Different symbols represent different timepoints (● = 1-day- old and ▴ = 28-day-old). Blue symbols represent foals that remained healthy after intrabronchial infection, gray symbol the foal that developed subclinical pneumonia, and red symbols foals that had clinical pneumonia. Statistical difference is represented by * for P < 0.05, **** for P < 0.00001, and ns for not significant.

**Fig. 8:**
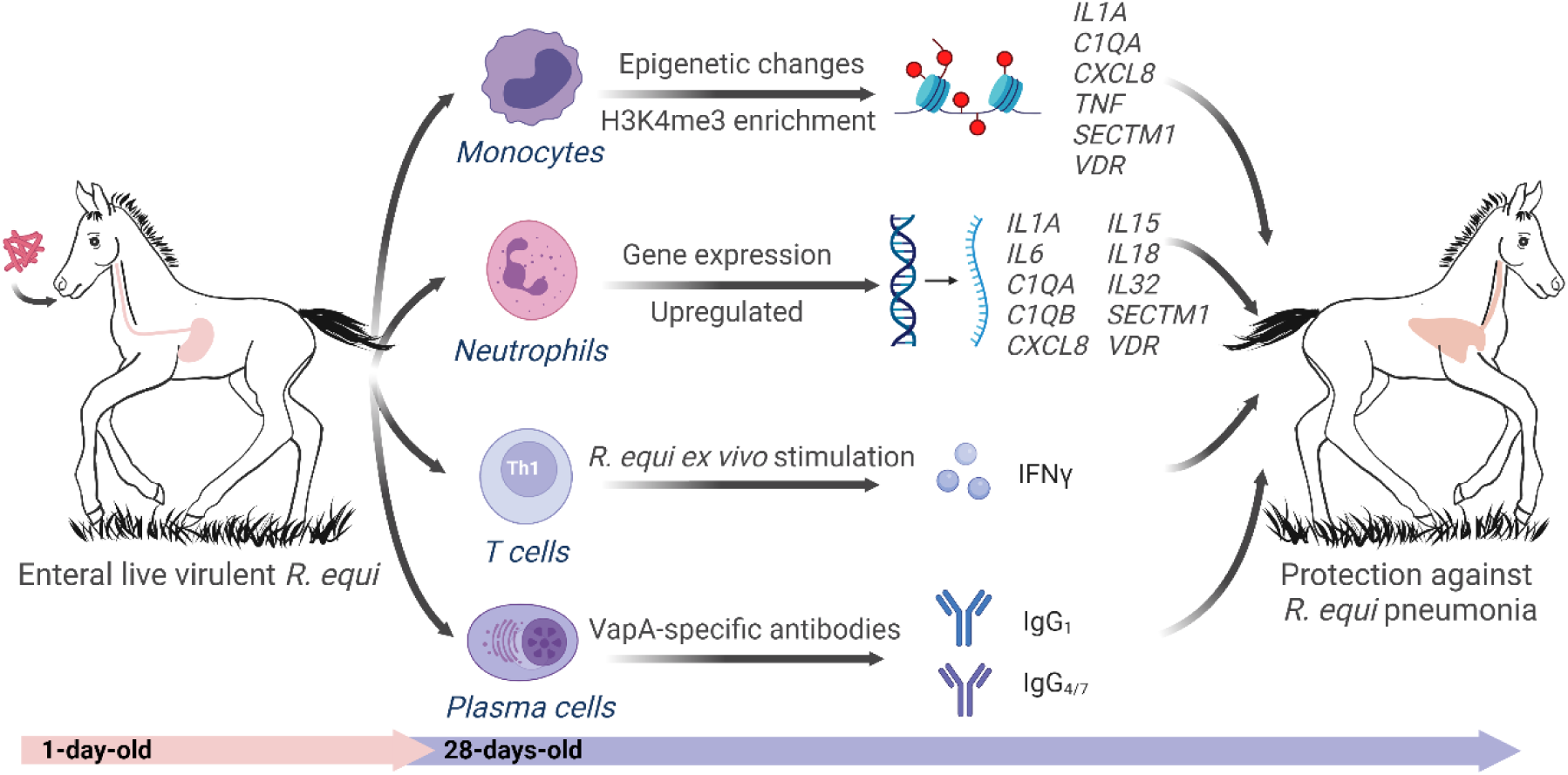
Summary of our results. Foals immunized enterally with *R. equi* at ages 1 and 3 days developed enhanced innate and adaptive immune responses at age 28 days. Innate cells of immunized foals had several important changes by age 28 days, including a modified epigenetic profile in monocytes and more than 1,000 differentially-expressed genes (DEG) in neutrophils. Adaptive immune responses to immunization included cell-mediated immunity (increased number of IFN-γ producing cells in PBMCs) as well as humoral responses (increased VapA-specific IgG_1_ and IgG_4/7_ activity in serum). Together, these enhanced innate and adaptive immune responses protected foals against intrabronchial challenge with *R. equi*.

## Discussion

Here we demonstrate that enteral immunization with *R. equi* in the first few days after birth generated protective immunity against pneumonia, suggesting the gut-lung axis is active in promoting immunogenicity in newborn foals (*43*), despite immaturity and naivety of their innate and adaptive immune systems (*2, 3, 17–21*). We also demonstrate for the first time that enteral immunization with *R. equi* enhanced innate immune responses, expanding knowledge from previous studies that focused only on induction of adaptive immune responses (*33, 34*). This is impactful because enhanced function of the innate immune system appears to be an important step in protecting foals against *R. equi*, presumably because innate immune cells directly kill bacteria (*44–46*), orchestrate the immune response (*47*), and facilitate adaptive immune response through more effective antigen presentation (*48, 49*).

To test our hypothesis that enteral immunization induces enhanced innate immune responses, we performed ChIP-qPCR in blood neutrophils and monocytes from foals, as well as transcriptome analysis in foal neutrophils. Our results indicate that enteral immunization with *R. equi* induced an innate immune profile similar to that previously observed with administration of BCG, a well-recognized inducer of trained immunity (*9, 41, 50, 51*). Notably, we found epigenetic changes in monocytes at the promoter regions of genes previously associated with trained immunity, such as *IL1B*, *IL32*, *CXCL8*, and *TNF* (*51–53*). Of note, *IL1A* might be especially important in horses in addition to *IL1B* (traditionally associated with trained immunity in other species) because we found both epigenetic changes in monocytes and increased gene expression of *IL1A* in neutrophils. Moreover, expression of the gene encoding the IL-1 receptor antagonist (*IL1RN*) was downregulated and *IL18*, a receptor agonist from the IL-1 family of cytokines, was upregulated in neutrophils from immunized foals, potentiating the IL-1 signaling pathway. The IL-1 family of cytokines control pro-inflammatory reactions, trained immunity induction, immunological defense against microbial infections, and the activation of T lymphocytes (*54*).

Additionally, we found epigenetic changes in other genes that encode proteins serving important immune functions, such as *C1QA*, *SECTM1*, and *VDR*. Complement component 1q (C1q) is part of the complement cascade and represents an important bridge between the innate and adaptive immune systems. Specifically, C1q deposition enhances antibody-mediated *R. equi* killing by equine neutrophils (*55*), and in people and mice, neutrophil production of C1q protects against sepsis by inducing resolution of inflammation and clearance of apoptotic neutrophils by macrophages (*56*). In immunized foals, we observed an enrichment of H3K4me3 in the *C1QA* promoter in monocytes as well as increased gene expression of *C1QA* and *C1QB* in neutrophils, suggesting this can be an important pathway for protecting foals against *R. equi* pneumonia. Similarly, *SECTM1*, a gene with important role as a costimulatory ligand for T-cell proliferation (*57, 58*) and chemoattractant for monocytes (*59*), was upregulated in neutrophils and exhibited H3K4me3 enrichment in monocytes from immunized foals. The specific role of *SECTM1* in the immune response to enteral *R. equi* merits further evaluation. Moreover, we observed in immunized foals that *VDR* gene, encoding vitamin D receptor (VDR), was upregulated in neutrophils as well as the H3K4me3 was enriched in promoter regions of *VDR* in monocytes. Expression of VDR is necessary for neutrophils to respond to vitamin D, which dampens excessive inflammation and apoptosis, and increases bacterial killing as observed for *Streptococcus pneumoniae* (*60*). Human neonatal neutrophils have decreased VDR expression and impaired anti-inflammatory activity of vitamin D (*61*). In foals, a recent preliminary study showed that vitamin D metabolism and VDR gene expression by alveolar macrophages is impacted by age (*62*). Deficiency of VDR expression in neonatal foals compared to adult horses, its role in susceptibility to pneumonia, and this increased expression following enteral immunization requires further elucidation.

Additionally, we observed that monocytes from immunized foals had H3K4me3 enrichment in the promoter region of *RPL30,* which is a transcriptionally active gene in most cell types and is commonly used as control for ChIP. In human monocytes, the H3K4me3 modification is acquired throughout the genome during development from neonate to adult, including in *RPL30* and other immunologically important genes such as *CCR2*, *IL1B*, and *TNF* (*63*). Therefore, our results suggest that enteral *R. equi* prompted the deposition of H3K4me3 and possibly accelerated the maturation of monocytes from a newborn to an adult phenotype (*63*).

Monocytes and neutrophils originate from the same myeloid progenitor cells, yet immunization induced H3K4me3 enrichment of selected genes in monocytes but not in neutrophils. We observed, however, that several of these genes were differentially-expressed in neutrophils from immunized foals. We believe that this discrepancy might be due to the heterogeneity of the neutrophil population. Previously, single-cell RNA-seq data indicated that even when only a small subset of the hematopoietic stem and progenitor cells (HSPCs) and circulating immune cells (i.e., neutrophils and monocytes) are trained, they can generate a protective phenotype (*36*). Single-cell transcriptomics are needed to determine whether a similar effect occurs in neutrophils, monocytes, and HSPCs from immunized foals.

We observed in both neutrophils and monocytes that the promoter regions of several genes (*IL32*, *C1QA*, *CCL8*, *SECTM1*, and *VDR*) had enrichment of both the H3K4me3 (associated with active transcription) and H3K27me3 (associated with transcriptional repression); this unique signature composed of both chromatin modifications classifies these regions as bivalent domains (*64*). Bivalent domains have been proposed to poise genes with critical or developmental roles for rapid expression upon differentiation, quickly allowing activation while maintaining repression in the absence of activation signal (*64*). Some bivalently marked genes can lose H3K27me3 and become expressed, or lose H3K4me3 and become silenced (*65*). It is unclear what controls the fate of bivalent genes. In human HSPCs, about 24% of bivalent domains remained after differentiation of the HSPCs into erythrocyte precursor cells (*65*). Here we showed that enteral immunization induced upregulation of genes marked with bivalent domains. In particular, 3 genes (*C1QA*, *CCL8*, and *SECTM1*) marked with bivalent domain were within the 6 highest fold-change in expression in the immunized group with > 34-fold change compared to controls. This finding suggests that genes marked with a bivalent domain in foal neutrophils can be highly expressed and that – for some genes – their expression was associated with protection against *R. equi* pneumonia.

The gut microbiota mediates lung mucosal immunity though the gut-lung axis, playing a protective role in respiratory infections (*30–32*). Absence or depletion of the gut microbiota in mice impairs immune responses and results in increased susceptibility to bacterial respiratory infection (*30–32*). The gut-lung axis also mediates the training of alveolar macrophages by parenteral administration of BCG vaccine through an intestinal microbiota-mediated pathway, which induces innate memory of distal mucosal-resident innate cells, and results in protection against pulmonary tuberculosis (*50*). In neonatal mice, exposure to intestinal commensal bacteria immediately after birth promotes a robust host defense against bacterial pneumonia (*43*). *R. equi* is commonly found as part of the intestinal microbiota of healthy horses (*66*). Therefore, we speculate that the gut-lung axis is involved in enhancing immune responses and protecting against pneumonia induced by enteral *R. equi*. Additionally, the natural resistance to *R. equi* pneumonia observed in some foals (*16, 24, 67*) could be due to ingestion of fecal *R. equi* present in the farm environment because coprophagia is common in foals (*68*). These ingested *R. equi* could stimulate maturation of the innate immune system.

It has been hypothesized that the inability of foals to develop a Th1 response and lower expression of IFN-γ at birth explains their heightened susceptibility to *R. equi* (*19, 20, 69*). Th1 deficiency could result from inefficient antigen presentation by immature antigen presenting cells that have lower expression relative to adult horses of CD1 (an MHC class I homologue), MHC class II, or co-stimulatory factors and cytokines (*2–5*). Previously, trained immunity has been suggested to increase MHC II expression and improve antigen presentation (*36*), and recent evidence shows the potential for neutrophils to participate in the activation of T-cells (*70, 71*). Here, we found upregulation of several genes associated with antigen processing and presentation in neutrophils from immunized foals and a *R. equi*- specific Th1 immune response. In the humoral immune responses, the antibody subisotype IgG_1_, associated with a Th1-biased immune response (*72*), was proportionally higher than that of IgG_4/7_ in immunized foals compared to our positive control serum from *R. equi*- hyperimmunized horses. IgG_1_ has also been associated with protection against *R. equi* pneumonia (*72*) and increased antibody-mediated bacterial killing by neutrophils (*55*). These results suggest stimulation of innate immune cells by enteral *R. equi* immunization may facilitate antigen presentation and allow foals to develop improved Th1 immune responses. Moreover, the increased IFN-γ production by *R. equi*-stimulated PBMCs and the upregulation of several IFN-γ-induced genes that we observed suggest that our immunization protocol reversed foal IFN-γ deficiency.

A previous vaccine trial using gavage of inactivated *R. equi* failed to protect foals against pneumonia, despite induction of a *R. equi*-specific cell-mediated immune response and local IgA (*24*). Immunization in mice with live virulent *R. equi*, but not killed or avirulent *R. equi*, also elicits protective immunity against *R. equi* (*73*). For other bacteria known to induce trained immunity, such as BCG vaccine strains and *Shigella*, inactivation weakened or abrogated the training of innate immune cells (*50, 74*). Together, our results indicated that *R. equi* needs to be alive to stimulate protective innate immune response.

Although these results are exciting, our study has limitations. First, we were not able to definitively determine whether the enhanced innate immune response observed in immunized foals was due to either memory features of hematopoietic progenitor cells (trained immunity) or continuous *R. equi* stimulation in the gut (constant priming of innate immune responses) because *R. equi* survives for weeks in the gut after enteral administration (*68*). Both hypotheses need to be considered because there are similarities between immunization with enteral *R. equi* and BCG vaccine, which is a well-recognized inducer of trained immunity (*9, 41, 50, 51*). These include similarities between the 2 microorganisms, as well as the innate immune responses to both enteral *R. equi* and the BCG vaccine. Also, gut bacteria can direct innate immune cell development by promoting hematopoiesis that limits bacterial infection (*75*). To further determine whether gut stimulation with *R. equi* induced trained immunity, *R. equi* must be removed after the prime immunization followed by a resting period in the absence of *R. equi* before intrabronchial infection. This poses a challenge given the ubiquitous nature of *R. equi* in the environment on horse farms. Second, we did not examine the mechanisms and intestinal cells directly interacting with *R. equi* in the gut lumen or the indirect stimulation of HSPCs in the bone marrow. Third, we did not explore the possible induction of protection against a heterologous microbe that would further substantiate enhanced innate immunity by cross-protection. Our study, however, is the first step to demonstrate the important role of the innate immune system in the protection induced by *R. equi* enteral immunization, and potential ability of *R. equi* to induce enhanced innate immunity.

Experience with existing vaccines indicates that newborns can mount robust immune responses to some vaccines, particularly to live vaccines (*8*). We showed that enteral immunization with *R. equi* induces changes in the gene expression of neutrophils and the epigenetic profile of monocytes, as well as *R. equi*-specific IgG_1_, IgG_4/7_, and Th1 IFN-γ. Together, these changes were associated with protection against experimental challenge and development of clinical pneumonia. Our data support the hypothesis that enteral immunization accelerates the process of maturation of the newborn foal immune system, corroborating previous studies indicating that exposure to environmental microorganisms promotes maturation of innate immune cells (*21*). Collectively, our findings indicated that enteral immunization with *R. equi* enhances maturation and effectiveness of innate immune cells, which is associated with protecting foals against rhodococcal pneumonia.

## Materials and Methods

### Experimental design and subject details

We designed our experiments following ARRIVE guidelines, and the Texas A&M University Institutional Animal Care and Use Committee reviewed and approved all procedures for this study (protocol number IACUC 2020-0273). We used a total of 11 healthy outbred Quarter Horse foals. Sample size calculations were performed based on the following assumptions: binomial distribution; statistical power = 80%; significance level of 5%, that all foals immunized with enteral *R. equi* would be protected (based on prior reports) (*27–29*); and, that 80% of control foals will develop pneumonia (based on prior reports) (*24*). BPS and SKK assigned pregnant mares randomly to the Control and Immunized groups before birth, with the sex unknown. At birth we obtained Control (n=6; 4 males and 2 females) and Immunized (n=5; 4 males and 1 female) foals (Fig. 1A), which were healthy and had transfer of passive immunity, as assessed by a commercially available qualitative immunoassay for serum concentration of total IgG (SNAP test; IDEXX, Inc., Westbrook, USA) ≈24 hours after birth. Foals exhibited age-appropriate results of complete blood count (CBC) at ages 1 and 28 days (Fig. S6). We monitored foals at least twice daily for 12 weeks.

### Bacteria preparation

We prepared fresh culture of *R. equi* strain ATCC 33701^+^ (ATCC, Manassas, Virginia) for enteral administration (*28*). We grew *R. equi* overnight in brain-heart infusion broth (BHI; BD Biosciences, St. Louis, MN) at 37 °C, washed twice with Phosphate-buffered saline (PBS; Gibco, Waltham, MA), and adjusted the concentration by optical density. For intrabronchial infection, frozen stock of *R. equi* strain EIDL-5331 obtained from a Texas foal with clinical pneumonia was be used (*24, 67*).

### Enteral administration of *R. equi* to foals

We randomly assigned individual foals at ages 1 and 3 days to receive enteral administration via nasogastric tube (*33, 76*) of either saline (Control Group; NaCl 0.9%; 150 ml; n=6) or *R. equi* (Immunized Group; 2 × 10^10^ *R. equi* in 150 ml of NaCl 0.9%; n=5). We maintained foals and their respective dams at a single facility in stalls for 7 days after birth, and then housed in paddocks until age 28 days. After challenge, foals were housed in stalls for 10 days and then moved to paddocks where they were maintained until the end of the study.

### Experimental infection in foals

Before experimental infection, we documented the absence of pre-existing lung lesions by thoracic ultrasonography (TUS). We infected foals with *R. equi* at age 28 days using our non-lethal model (Fig. 1A) (*24, 46, 67, 68*). Briefly, we sedated foals using intravenous (IV) injection of xylazine (0.2 mg/kg) and butorphanol tartrate (0.04 mg/kg). An aseptically- prepared videoendoscope with an outer diameter of 9-mm was inserted via the nares into the trachea and passed to the bifurcation of the main-stem bronchi. We inserted a disposable sterile endoscopy catheter (Endoscopy Support Services, Inc.) to transendoscopically administer a 50-ml suspension containing approximately 2 × 10^6^ total viable virulent *R. equi*, with 25 ml each infused into the right and left mainstem bronchus.

### Clinical monitoring

We performed twice daily physical examination, which included recording the temperature, respiratory and heart rate from birth. Beginning on the day of infection through age 84 days, we recorded the following: 1) rectal temperature, heart rate, and respiratory rate (Fig. S7); 2) tracheal and thoracic auscultation for abnormal airway sounds; 3) cough; 4) increased respiratory effort; and, 5) attitude, including posture and frequency of suckling. The pre- determined case-definition of pneumonia was the presence of ≥ 3 of the following clinical findings: 1) cough at rest; 2) rectal temperature > 39.4 °C; 3) respiratory rate ≥ 60 and increased respiratory effort at rest; 4) abnormal lung sounds; 5) abnormal tracheal sounds (rattle); and, 6) lethargy. We performed weekly CBC and TUS to identify evidence of peripheral pulmonary consolidation or abscess formation consistent with *R. equi* pneumonia. AIB and NDC determined the pneumonia status of each foal and were masked to the immunization groups until the end of the study. At the onset of clinical signs of pneumonia, we performed the following tests: CBC, TUS, and a transendoscopic tracheo- bronchial aspirate (T-TBA). For the T-TBA, we sedated foals with xylazine and butorphanol at previously described doses. We used a sterile triple-guarded tubing system and sterile 0.9% physiological saline solution to collect the T-TBA aseptically for cytologic examination and microbiologic culture. We recorded the presence of either cytologic evidence of septic inflammation or Gram-positive pleomorphic coccobacilli and results of microbiologic culture of *R. equi*.

Following diagnosis of rhodococcal pneumonia, we treated pneumonic foals with clarithromycin (7.5 mg/kg; by mouth; q 12 h) and rifampin (5 mg/kg; by mouth; q 12 h) until both clinical signs and TUS lesions had resolved. If needed to treat fever, we used flunixin meglumine (0.6 to 1.1 mg/kg; by mouth; q 12 h). At age 84 days, foals were completely recovered and a second T-TBA was performed to confirm absence of cytologic sepsis and negative culture for *R. equi*.

### *R. equi* detection on blood

We collected 3 ml of blood from the jugular vein into an ethylenediaminetetraacetic acid (EDTA) vacuum tube at ages 1, 2, 4, 6, 14, 28, 42, and 56 days of age for detection of *R. equi* in blood (Fig. 1A). For *R. equi* bacterial enrichment, we added 1 ml of blood into 50 ml of *R. equi*-selective NANAT broth and incubated at 120 rpm at 37 °C for 48 h. Next, we plated 20 µl of NANAT broth culture in BHI agar and kept at 37 °C for 48 h. We extracted the DNA from colonies with morphology compatible with *R. equi* using DNeasy Blood & Tissue Kit (Qiagen, Hilder, Germany). We determined DNA quality and quantity via spectrophotometer (Nanodrop 1000, Thermo Fisher Scientific, Waltham, MA) before storage at -20 °C. We performed real-time PCR using 10 µl total volume reaction, TaqMan™ Fast Advanced Master Mix (Thermo Fisher Scientific, Waltham, MA), and *vapA* specific TaqMan™ primer and probe (Forward: GAGCAGCAGTACGACGTTCA; Reverse: GGCCCGAATACGTGAAACCT; Reporter: CAGCGCGGTCGTCTAC) in the instrument QuantStudio 6 Flex Real-Time PCR System (Applied Biosystems).

### Neutrophil, PBMC, and monocyte isolation

We collected blood from the jugular vein into heparinized vacuum tubes from all foals at ages 1 (before enteral immunization) and 28 days (before intrabronchial challenge) (Fig. 1A). We separated neutrophils, monocytes, and peripheral blood mononuclear cells (PBMCs) using a double layer density separation (Histopaque; densities = 1.077 and 1.191; Sigma-Aldrich, Gillingham, United Kingdom) (*21, 77*) and washed with cold Hanks’ Balanced Salt Solution (HBSS; Gibco, Waltham, MA). We lysed remaining red cells in the neutrophil fraction with 0.84% NH_4_Cl incubating for 3 min. We washed neutrophils and PBMCs suspensions 2 × with cold HBSS and counted using an automated cell counter (Cellometer Auto T4, Nexcelcom, MA); we used PBMCs either for ELISpot or for monocyte isolation by selective adherence as previously described (*78*). We resuspended neutrophils and monocytes in Roswell Park Memorial Institute 1640 medium (RPMI; Lonza, Walkersville, MD) supplemented with 10% horse serum to a concentration of 3.3 × 10^6^ cells/ml and immediately fixed for ChIP (see below). For gene expression and mRNA sequencing, we lysed 5 × 10^6^ pelleted neutrophils with 300 µl of TRIzol® Reagent (Life Technologies, Carlsbad, CA) and stored at -80 °C. We failed to isolate cells from one foal in the control group at age 28 days because of the presence of low-density neutrophils that interfere with the density gradient separation.

### Gene expression of neutrophils

We isolated total RNA from samples of neutrophils frozen in TRIzol® Reagent (Life Technologies™, Carlsband, CA) using Direct-zol RNAeasy kit (Zymogen) (*78*). We determined RNA quality and quantity via spectrophotometer (Nanodrop 1000, Thermo Fisher Scientific, Waltham, MA). We used 200 ng of RNA for cDNA synthesis using RT2 First Strand Kit (Qiagen Sciences, Germantown, MD) according to manufacturer’s instructions. We performed gene expression assays for 8 selected genes (*TNF, IL1A, IL6, IL1RN, ATG5, CXCL8, IL32,* and *STAT4*) using 1 ng of cDNA, TaqMan Gene Expression Assays (ThermoFisher, Waltham, MA) (Table 1), and TaqMan Fast Advanced Master Mix (Thermo Fisher Scientific, Waltham, MA) according to manufacturer’s instructions. We performed reactions in 384-well plate in triplicates. We chose *B2M* as the endogenous control to normalize fold-changes in transcripts levels, QuantStudio 6 Flex Real-Time PCR System (Applied Biosystems) for relative quantification, and the relative quantification 2^-ΔΔCt^ formula for fold-changes analyses.

**Table 1:** Primers used in neutrophil gene expression.

| Gene | Gene encodes | Function | Assay ID |
| --- | --- | --- | --- |
| *B2M* | Beta-2-Microglobulin | Housekeeping | Ec03468699_m1 |
| *TNF* | Tumor Necrosis Factor alpha | Pro-inflammatory | Ec03467871_m1 |
| *IL1A* | Interleukin 1 alpha | Pro-inflammatory | Ec03468697_m1 |
| *IL1RN* | Interleukin 1 Receptor Antagonist | Block IL1R (anti-inflammatory) | Ec03468814_m1 |
| *ATG5* | Autophagy Related 5 | Autophagy pathway | Ec07051187_m1 |
| *CXCL8* | C-X-C Motif Chemokine Ligand 8 | Chemotactic factor | Ec03468860_m1 |
| *IL-32* | Interleukin 32 | Pro-inflammatory, trained immunity | Ec07065202_g1 |
| *STAT4* | Signal Transducer and Activator of Transcription 4 | Antimicrobial immunity | Ec06959089_m1 |
| *IL6* | Interleukin 6 | Pro-inflammatory | Ec03468678_m1 |

### RNA sequencing

We isolated total RNA from foal neutrophil at ages 1 and 28 after birth as described above. Texas A&M Institute for Genome Sciences and Society (College Station, TX, USA) constructed sequencing libraries for each sample using a standard protocol (Illumina, San Diego, CA). RNA-Seq library quality was assessed with the Cutadapt package and mapped to the equine genome using the HISTAT2 package. We performed pathway analysis using IPA (Qiagen).

### ChIP-qPCR

To identify epigenetic changes on foal neutrophils and monocytes at ages 1 and 28 days, we used ChIP-qPCR targeting the promoter regions of 12 genes. We fixed approximately 1.6 × 10_7_ neutrophils or 1 × 10^6^ monocytes by adding paraformaldehyde 16% (Electron Microscope Sciences, Hatfield, PA) to a final concentration of 1% and incubating at room temperature for 10 min, inverting occasionally to mix. We stopped crosslinking by adding 10× of glycine solution (Cell Signaling Technology, Danvers, MA) to 1× concentration and incubating for 5 min inverting, occasionally to mix. Then, we washed cells 3× with cold PBS, resuspended in 10 ml of cold PBS, and kept at 4 °C for up to 10 days until chromatin fragmentation using SimpleChIP® Plus Sonication Chromatin IP Kit (Cell Signaling Technology, Danvers, MA). Briefly, we pelleted each sample (either neutrophils or monocytes), resuspended them in cell lysis buffer, and incubated for 10 min on ice. Then, we washed cells with cell lysis buffer, incubated them for 5 min, pelleted, resuspended in nuclear lysis buffer, and incubated for 10 min on ice. We transferred cells into 1.5 ml sonication tubes (200 µL/tube) and sonicated 30 sec on/30 sec off for 6 cycles at 4 °C using Bioruptor® Pico (Diagenode, Denville, NJ). We centrifuged fragmented chromatin at 20,800 g for 10 min at 4 °C, collected supernatants, and froze at -80 °C until immunoprecipitation (IP). To evaluate chromatin fragments, we incubated a 40 µL aliquot of each fragmented chromatin with RNAse A at 0.13 mg/ml at 37 °C for 30 min, and then reversed cross-link with proteinase K at 0.25 mg/ml at 65 °C for 2 h. Next, we column purified DNA and measured DNA concentration and quality via spectrophotometer (Nanodrop 1000, Thermo Fisher Scientific, Waltham, MA). We electrophoresed 10 µL of fragmented chromatin on 1.2% agarose gel with ethidium bromide and recorded the size of chromatin fragments to be about 150 - 300 bp. We performed ChIP using fragmented chromatin for each cell type (either 1 µg of monocytes or 6 µg for neutrophils), ChIP buffer with protease inhibitor cocktail, and rabbit mAb against specific targets for immune precipitation (IP; i.e., antibodies against histone H3, H3K4me3, H3K27me3, or normal IgG). Before adding mAb, we saved a 2% aliquot to be used for qPCR normalization. For IP, tubes with mAb were incubated on a rotator at 4 °C overnight on a rotator (Model 13916- 822, VWR, United States). These mAb included anti-histone H3 (1:50; clone D2B12; #4620; Cell Signaling Technology, Danvers, MA), anti-H3K4me3 (1:50; clone C42D8; #9751; Cell Signaling Technology, Danvers, MA), anti-H3K27me3 (1:50; clone C36B11; #9733; Cell Signaling Technology, Danvers, MA), or normal IgG (1:50; clone C2C12; #2729; Cell Signaling Technology, Danvers, MA). We precipitated the endogenous DNA-protein complex by adding 30 µl of ChIP-Grade Protein G Magnetic Beads, incubating on a rotator for 2 h at 4 °C, washing 2× with cold low-salt solution, washing 1× with high-salt solution, and eluting at 65 °C for 40 min. We decross-linked ChIP DNA and column purified as described above, then stored at -20 °C for future analysis by ChIP-qPCR.

We performed ChIP-qPCR using SimpleChIP® Universal qPCR Master Mix (Cell Signaling Technology, Danvers, MA), primers designed for the promoter region (0 – 1.5 kb upstream) of specific genes (Table 2), and ChIP-DNA (diluted 1:4). We performed reaction in triplicates with 10 µl total volume using QuantStudio 6 Flex Real-Time PCR System (Applied Biosystems). We normalized qPCR signal from ChIP DNA by 2% DNA input using the formula % of input = 100*2ᶺ(Ct [input]-Log_2_(50) – (Ct [ChIP]).

**Table 2:**
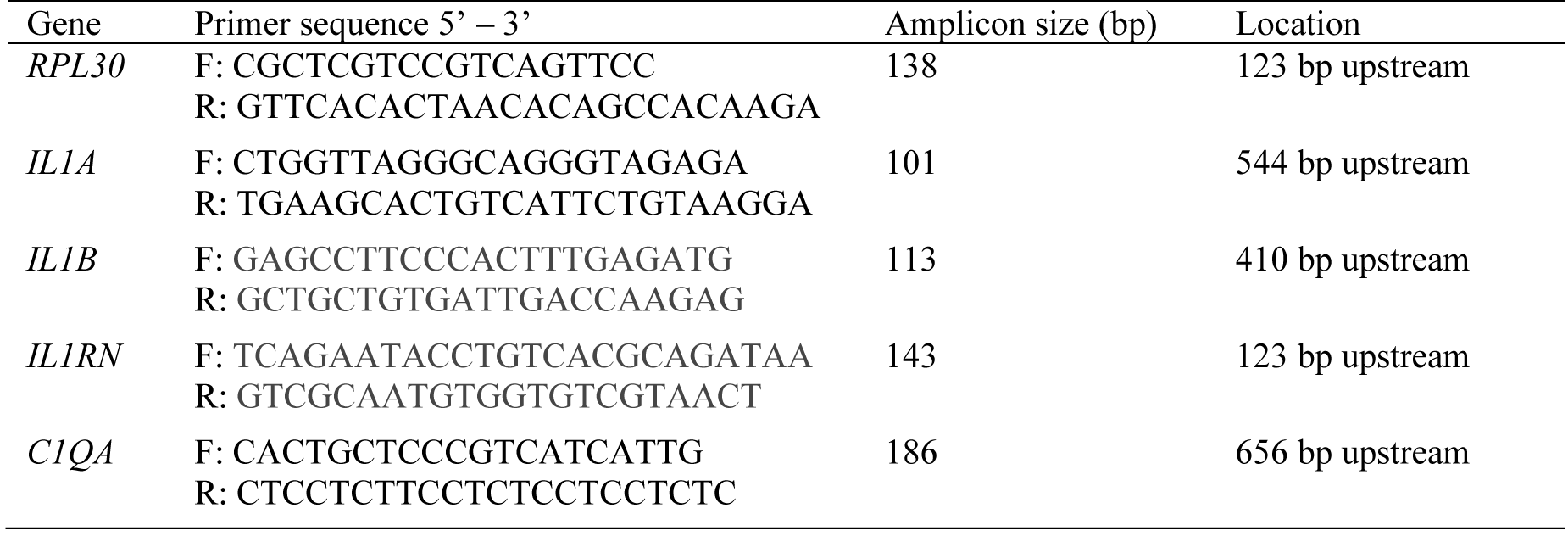
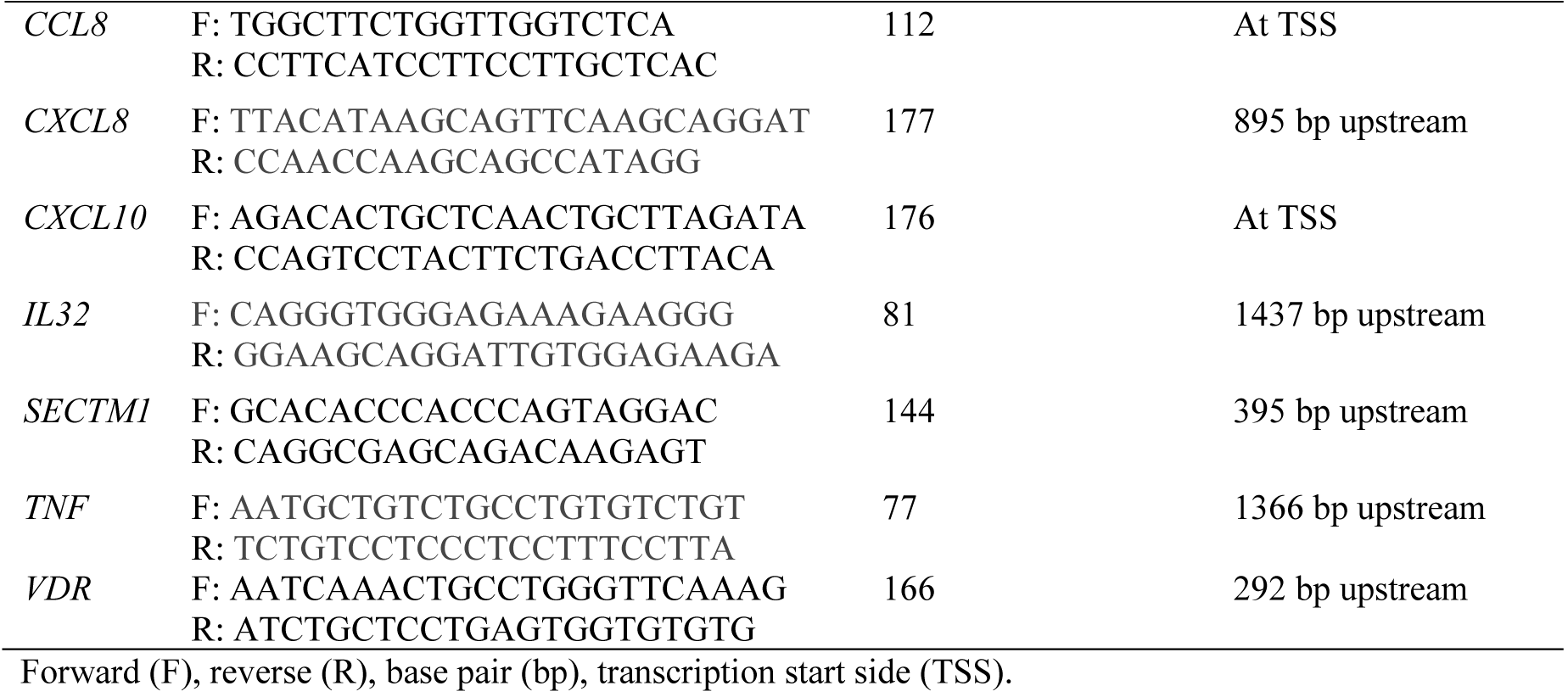
Primers used for ChIP-qPCR.

### Immunoglobulin ELISA

We tested serum samples collected at ages 1 and 28 days by enzyme-linked immunoassay (ELISA) for relative activities of antibodies against VapA protein of *R. equi*. We coated ELISA 96-well plates (Maxisorp, Thermo Scientific, Rochester, NY, USA) with 100 μl of 1 μg/ml purified recombinant VapA (rVapA) produced according to (*79*) diluted in sensitization buffer (0.04 M PO_4_, pH 7.2) overnight at 4°C (*80*). We washed plates 6 × with PBS containing 0.05% Tween-20 and blocked with 150 μl of PBS containing 1% skim milk for 1 hour at 37°C. We diluted foal serum samples in incubation buffer (PBS with 1% skim milk and 0.05% Tween-20) to 1:20 for IgG_1_ and 1:80 for IgG_4/7_ detection. We used *R. equi* hyperimmune serum (RE HIS; MG Biologics, Ames, IA) as positive control diluted to 1:160 for IgG_1_ and 1:10,240 for IgG_4/7_ detection. We added diluted serum samples (100 μl) in duplicate to wells of the ELISA plate and incubated for 1 hour at 37°C. We washed plates 6 ×, then added 100 μl per well of horseradish peroxidase (HRP) conjugated detection antibodies against either horse IgG_1_ (A70-124P 16; Bethyl Laboratories, Montgomery, TX, diluted at 1:40,000) or horse IgG_4/7_ (A70-123P 14; Bethyl Laboratories, Montgomery, TX, diluted at 1:80,000). We incubated plates for 1 hour at room temperature and then washed 6 ×. We added KPL SureBlue Reserve™ TMB Microwell Peroxidase Substrate One Component (100 μl; SeraCare, Gaithersburg, MD, USA) to the wells and incubated for 20 minutes. We stopped the reaction by adding 50 μl of 0.18 M sulfuric acid solution to the wells. We determined optical densities at a wavelength of 450 nm using a microplate reader (Synergy 2; BioTek, Winooski, VT). We calculated the relative antibody activity (OD ratio) by dividing the individual sample mean of OD values by the positive control mean from the same plate.

### ELISpot

To evaluating *R. equi*-specific Th1 response at age 28 days, we conducted *ex vivo* ELISpot assay using equine interferon-γ (IFN-γ) ELISpot kit (Mabtech, Inc.; Cincinnati, Ohio) according to manufacturer’s directions. First, we added 15 µl of 35% EtOH into ELISpot plates (S2EM004M99; Sigma-Aldrich, St. Louis, MO), washed 5 × with sterile water, coated with capture antibody (MT166; 15 µg/ml in PBS), and incubated overnight at 4 °C. We washed wells 5 × and blocked with cell culture medium (RPMI 1640 medium supplemented with 10% horse serum) for 30 min. Next, we added stimuli to designated wells in triplicate per condition. Our positive control was 0.5 µg of concanavalin A, negative was control media only, and sample test was 0.25 µg of *R. equi* lysate. Then, we added 100,000 PBMCs isolated from 28-day-old foals (as described above) in each well and incubated for 2 days in CO_2_ incubator at 37 °C. We washed wells 5 × with PBS, added 0.05 µg of detection mAb (MT13-biotin) in 100 µl of PBS, and incubated for 2 h at room temperature. We washed wells 5 × with PBS, added streptavidin-ALP (diluted to 1:1,000 in PBS and 5% FBS), and incubated for 1 h at room temperature. We developed with BCIP/NBT-plus substrate for ELISpot and stopped the reaction by washing with running water when distinct spots emerged. When plate was dry, we read spots using an AID plate reader (Advanced Imaging Devices GmbH, Strassberg, Germany).

### Statistical analysis

We analyzed clinical data (development of clinical pneumonia) from 6 control and 5 immunized foals using a 1-sided Fisher’s exact test, based on prior evidence that enteral immunization protects foals against *R. equi* pneumonia (*27–29*). We analyzed the presence of pulmonary lesions detected by thoracic ultrasound and ELISpot data using t-test. We analyzed gene expression, epigenetic changes, and IgG ELISA data using mixed-effects modeling with multiple pairwise comparisons made using the method of Tukey. RNA-Seq library quality was assessed with the Cutadapt package and mapped to the equine genome using the HISTAT2 package. Differential gene expression between groups was assessed using the DESeq2 package and q-value < 0.05. Significance was set at P < 0.05.

## Acknowledgments

Authors would like to thank Wesley Brashear, Lauren Hastings, Addison Bush, and Sophia C. Ramirez-Cortez for technical help. Fig. 1A and 8 were made using BioRender.com. We thank Idexx® Laboratories for donation of SNAP Foal IgG Tests and MG Biologics for donation of *R. equi* hyperimmune serum used as control for ELISA. We also thank the Veterinary Medical Park (Veterinary Medicine & Biomedical Sciences) and the Equine Nutrition Research and Reproductive Teaching Center (Department of Animal Science) staff.

## Funding

Grayson Jockey Club Research Foundation (AIB, NDC)

Boehringer-Ingelheim Animal Health (Advancement in Equine Research Award; AIB, NDC)

Department of Large Animal Clinical Sciences (AIB, NDC)

Link Equine Research Endowment, Texas A&M University (NDC)

## Author contributions

Conceptualization: BPS, NDC, AIB

Methodology: BPS, HMCP, SKK, RML, JMB, MCG, NDC, AIB

Investigation: BPS, NDC, AIB Visualization: BPS, NDC, AIB Supervision: NDC, AIB Writing—original draft: BPS, NDC, AIB

Writing—review & editing: SKK, RML, JMB, HMCP, MCG, BPS, NDC, AIB

## Data and materials availability

All data are available in the main text or the supplementary materials.

