## supplementary materials for "Enteral immunization with live bacteria reprograms innate immune cells and protects neonatal foals from pneumonia"

**Supplementary Materials**
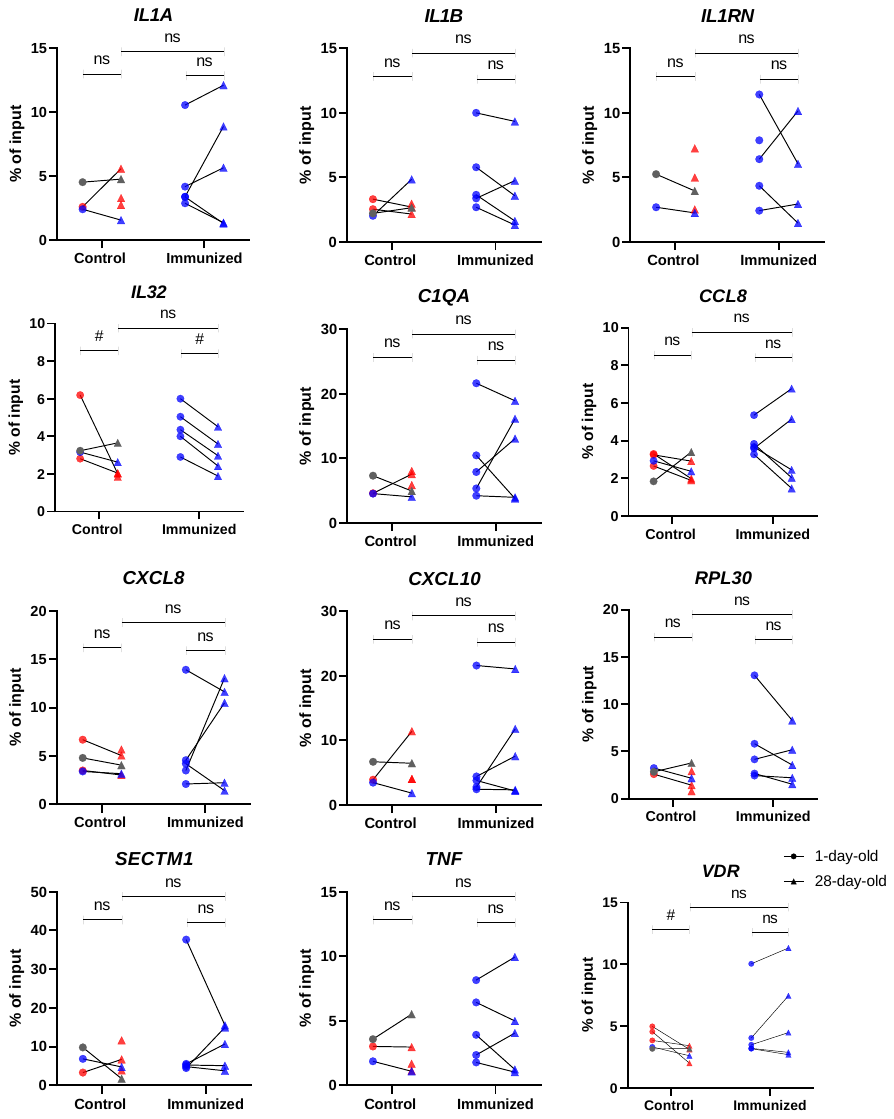

**Fig. S1:** **Enteral immunization did not induce H3K27me3 enrichment in promoter regions of genes in monocytes.** Blood monocytes were isolated from foals at 1- and 28-days of age. Chromatin-associated DNA was immunoprecipitated with mAb to the activating histone modification H3K27me3. DNA was purified and qPCR performed using primers specific for the region proximal to transcriptional start site (TSS) of *IL1A*, *IL1B*, *IL1RN*, *IL32*, *C1QA*, *CCL8*, *CXCL8*, *CXCL10*, *RPL30*, *SECTM1*, *TNF*, and *VDR*. Data is shown as % of input (% of input = 100*2^(Ct [input]-Log2(50) – (Ct [ChIP]). Different symbols represent timepoints (● = 1-day-old and▲ = 28-day-old). Different symbols represent different timepoints (● = 1-day-old and ▲ = 28-day-old). Blue symbols represent monocytes isolated from foals that remained healthy after intrabronchial infection, gray symbol the foal that developed subclinical pneumonia, and red symbols foals that had clinical pneumonia. The lines connect timepoints from the same animal. Statistical difference is represented by * for P < 0.05, # for 0.1 > P > 0.05, and ns for not significant.

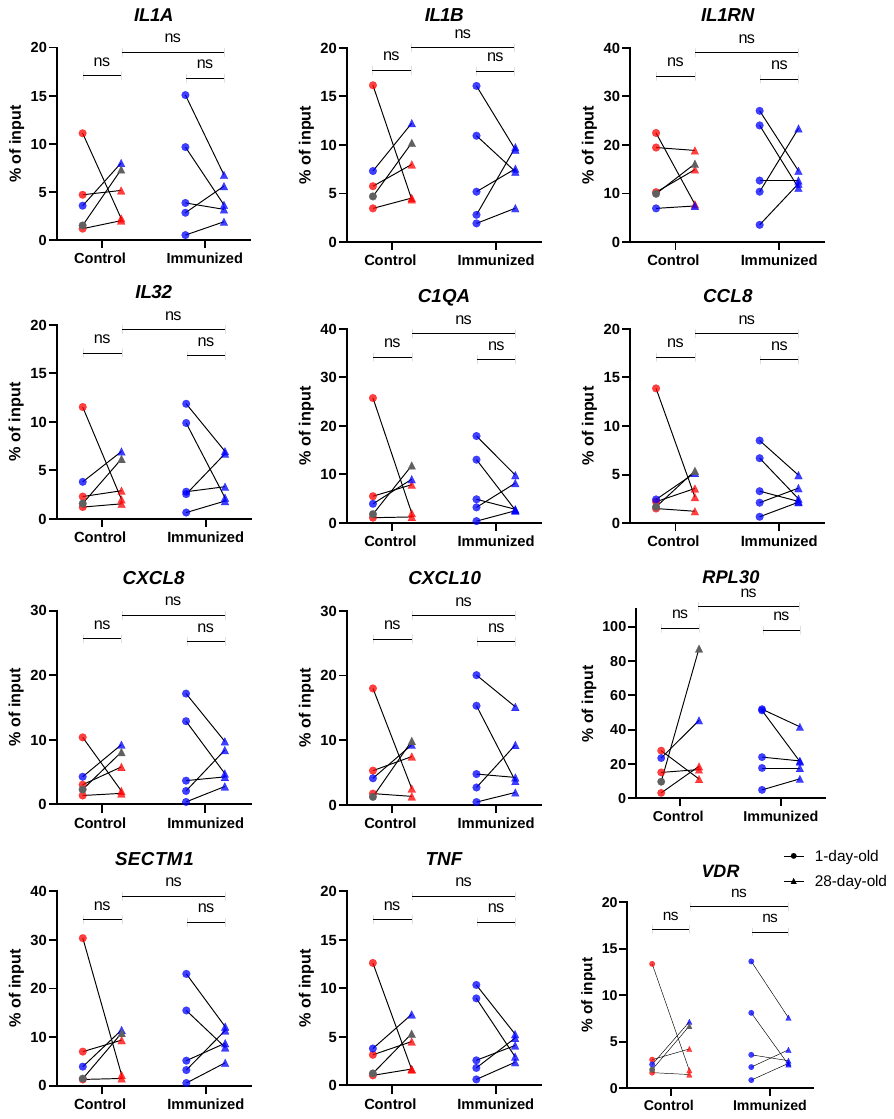

**Fig. S2:** **Enteral immunization did not induce H3K4me3 enrichment in promoter regions of genes in neutrophils.** Blood neutrophils were isolated from foals at 1- and 28-days of age. Chromatin-associated DNA was immunoprecipitated with mAb to the activating histone modification H3K4me3. DNA was purified and qPCR performed using primers specific for the region proximal to transcriptional start site (TSS) of *IL1A*, *IL1B*, *IL1RN*, *IL32*, *C1QA*, *CCL8*, *CXCL8*, *CXCL10*, *RPL30*, *SECTM1*, *TNF*, and *VDR*. Data is shown as % of input (% of input = 100*2^(Ct [input]-Log2(50) – (Ct [ChIP]). Different symbols represent timepoints (● = 1-day-old and▲ = 28-day-old). Different symbols represent different timepoints (● = 1-day-old and ▲ = 28-day-old). Blue symbols represent neutrophils isolated from foals that remained healthy after intrabronchial infection, gray symbol the foal that developed subclinical pneumonia, and red symbols foals that had clinical pneumonia. The lines connect timepoints from the same animal. No statistical difference is represented by ns.

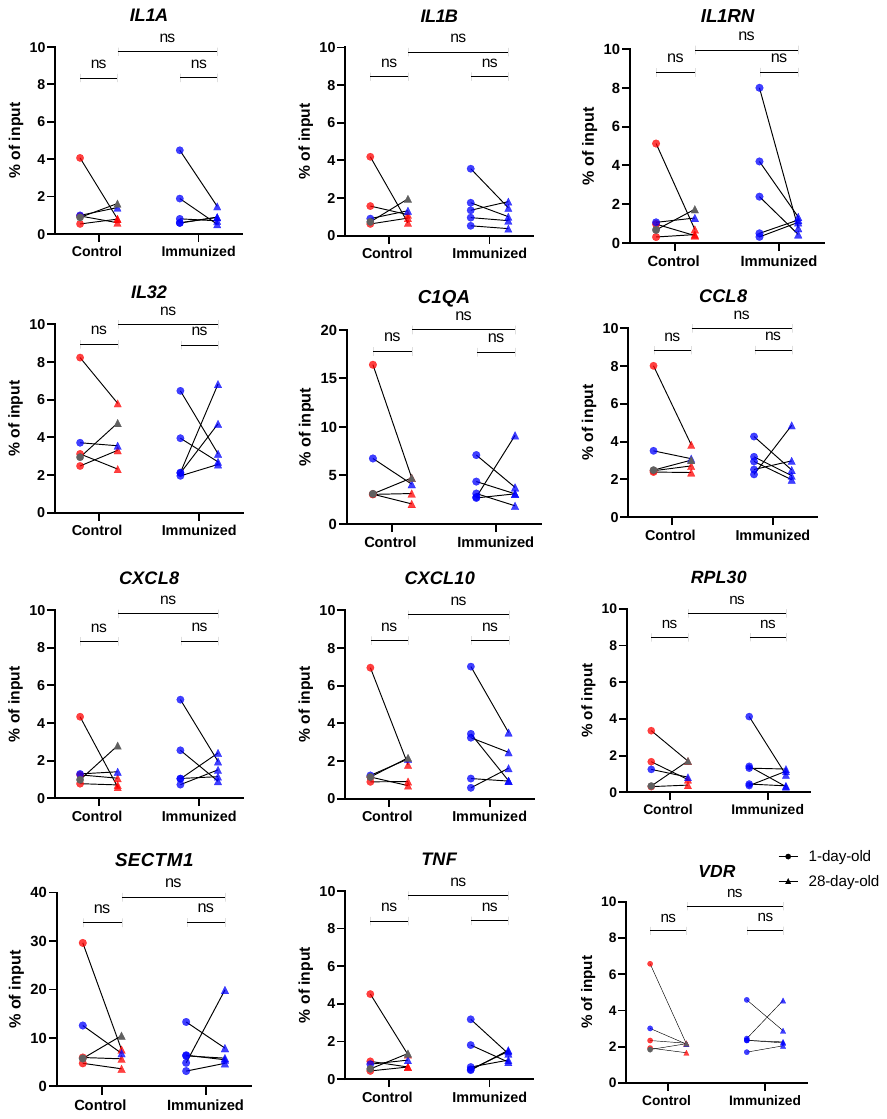

**Fig. S3:** **Enteral immunization did not induce H3K27me3 enrichment in promoter regions of genes in neutrophils.** Blood neutrophils were isolated from foals at 1- and 28-days of age. Chromatin-associated DNA was immunoprecipitated with mAb to the activating histone modification H3K27me3. DNA was purified and qPCR performed using primers specific for the region proximal to transcriptional start site (TSS) of *IL1A*, *IL1B*, *IL1RN*, *IL32*, *C1QA*, *CCL8*, *CXCL8*, *CXCL10*, *RPL30*, *SECTM1*, *TNF*, and *VDR*. Data is shown as % of input (% of input = 100*2^(Ct [input]-Log2(50) – (Ct [ChIP]). Different symbols represent timepoints (● = 1-day-old and▲ = 28-day-old). Different symbols represent different timepoints (● = 1-day-old and ▲ = 28-day-old). Blue symbols represent neutrophils isolated from foals that remained healthy after intrabronchial infection, gray symbol the foal that developed subclinical pneumonia, and red symbols foals that had clinical pneumonia. The lines connect timepoints from the same animal. No statistical difference is represented by ns.

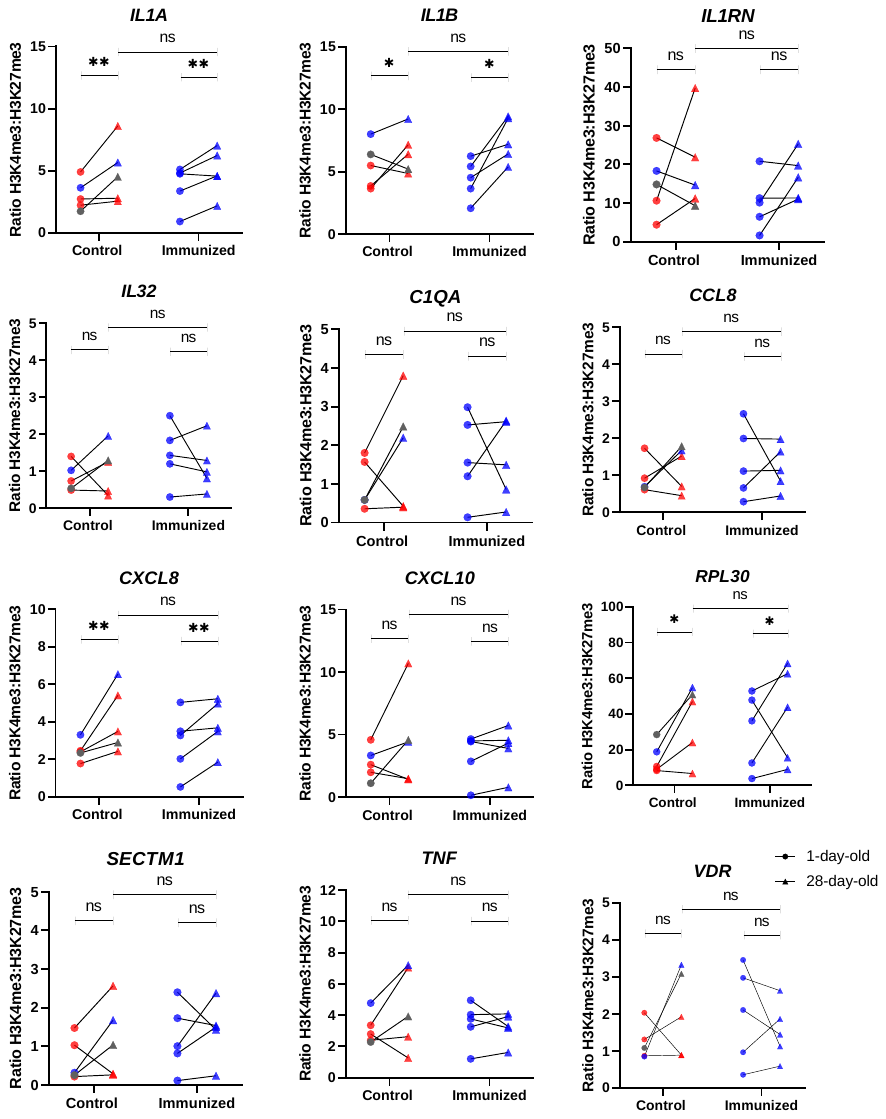

**Fig. S4:** **Enteral immunization did not alter H3K4me3:H3K27me3 ratio in promoter regions of genes in neutrophils.** Blood neutrophils were isolated from foals at 1- and 28-days of age. Chromatin-associated DNA was immunoprecipitated with monoclonal antibodies (mAbs) to the activating histone modification, H3K4me3, or the silencing histone modification, H3K27me3. DNA was purified and qPCR performed using primers specific for the region proximal to transcriptional start site (TSS) of *IL1A*, *IL1B*, *IL1RN*, *IL32*, *C1QA*, *CCL8*, *CXCL8*, *CXCL10*, *RPL30*, *SECTM1*, *TNF*, and *VDR*. Percentage of input was calculated as % of input = 100*2^(Ct [input]-Log2(50) – (Ct [ChIP]). Data show the ratio of % of input of H3K4me3 divided by % of input of H3K27me3. Different symbols represent timepoints (● = 1-day-old and▲ = 28-day-old). Different symbols represent different timepoints (● = 1-day-old and ▲ = 28-day-old). Blue symbols represent neutrophils isolated from foals that remained healthy after intrabronchial infection, gray symbol the foal that developed subclinical pneumonia, and red symbols foals that had clinical pneumonia. The lines connect timepoints from the same animal. Statistical difference is represented by ** for P < 0.01, * for P < 0.05, # for 0.1 > P > 0.05, and ns for not significant.

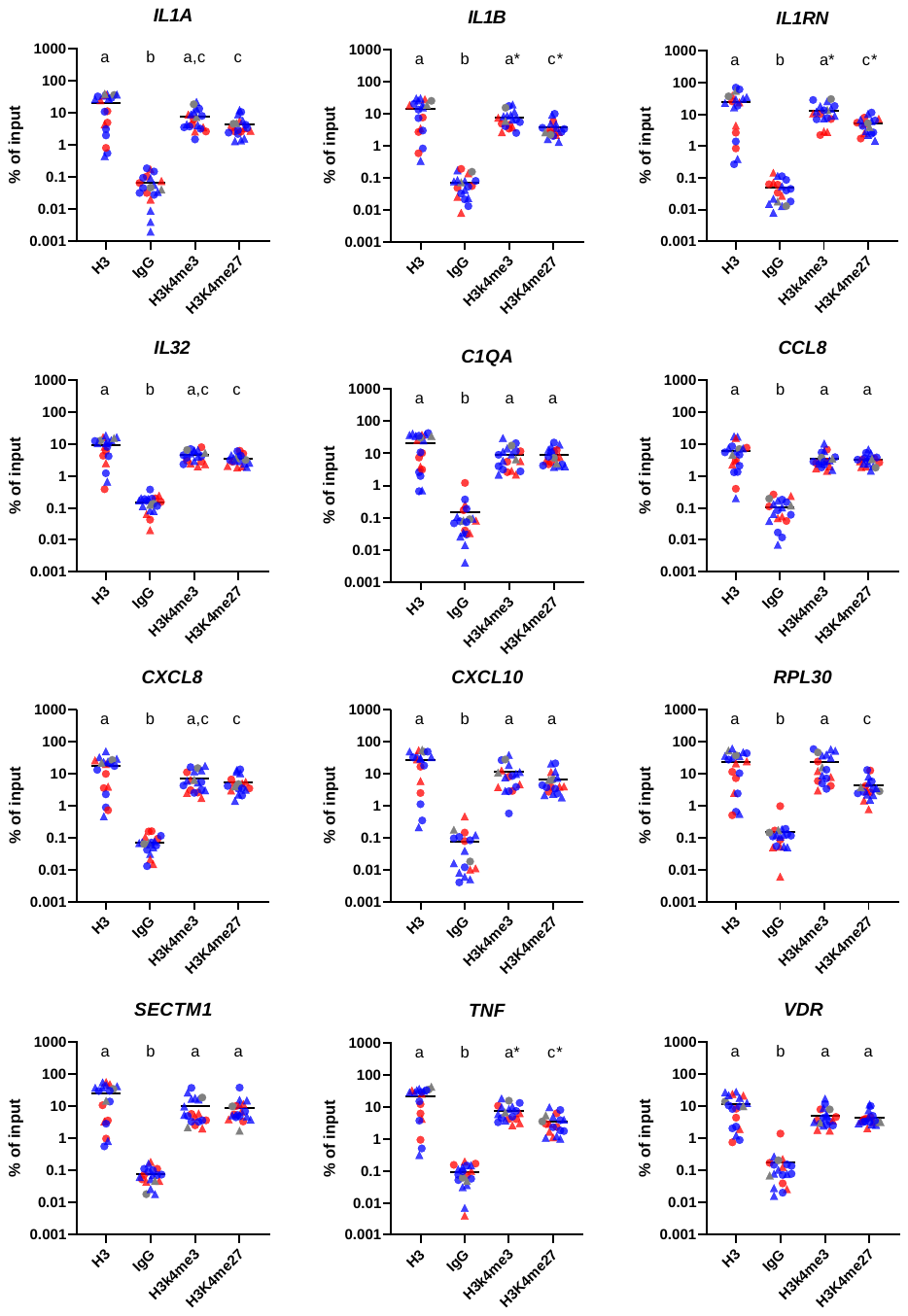

**Fig. S5:** **Monocytes from foals have a bivalent epigenetic signature for *IL1A*, *IL32*, *C1QA*, *CCL8*, *CXCL8*, *CXCL10*, *SECTM1*, and *VDR***. Blood monocytes were isolated from foals at 1- and 28-days of age. Chromatin-associated DNA was immunoprecipitated with monoclonal antibodies (mAbs) to the normal H3 histone (H3), normal IgG (mock control), activating histone modification (H3K4me3), or the silencing histone modification (H3K27me3). DNA was purified and qPCR performed using primers specific for the region proximal to transcriptional start site (TSS) of *IL1A*, *IL1B*, *IL1RN*, *IL32*, *C1QA*, *CCL8*, *CXCL8*, *CXCL10*, *RPL30*, *SECTM1*, *TNF*, and *VDR*. Percentage of input was calculated as % of input = 100*2^(Ct [input]-Log2(50) – (Ct [ChIP]). Bivalent domains were defines as having enrichment for both H3K4me3 and H3K27me3. Different symbols represent timepoints (● = 1-day-old and▲ = 28-day-old). Different symbols represent different timepoints (● = 1-day-old and ▲ = 28-day-old). Blue symbols represent monocytes isolated from foals that remained healthy after intrabronchial infection, gray symbol the foal that developed subclinical pneumonia, and red symbols foals that had clinical pneumonia. Statistical difference (P < 0.05) is represented by different letters (a, b, c, and d), and * for 0.1 > P > 0.05.

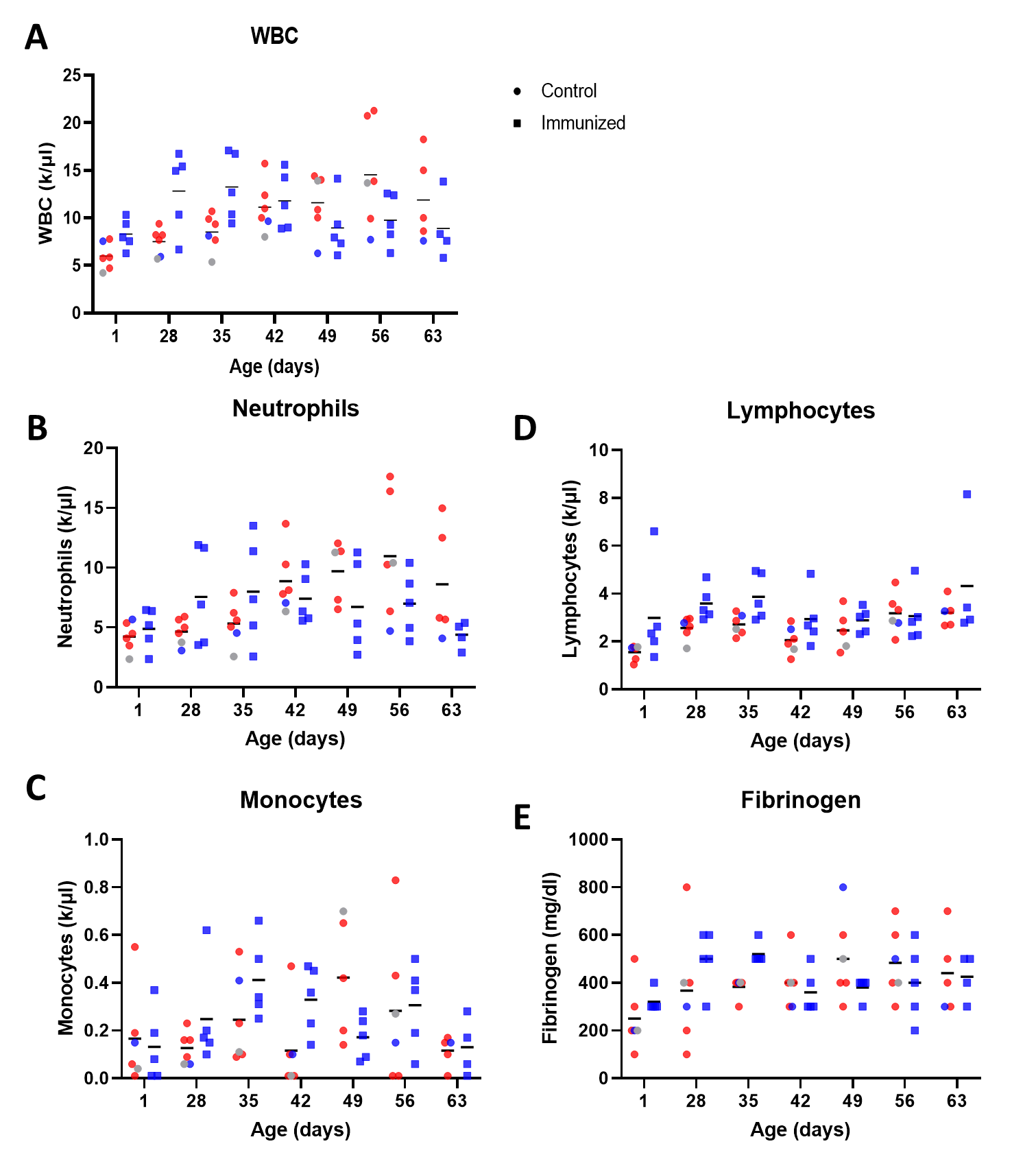

**Fig. S6:** **Complete blood cell count (CBC) results from birth to age 63 days.** Serial CBCs were performed by a reference laboratory. Total concentrations of leukocytes (white blood cells [WBC]; A), neutrophils (B), monocytes (C), lymphocytes (D), and fibrinogen quantification were reported from foal blood. Foals in the control group are represented by circles and immunized group by squares. Different symbols represent different timepoints (● = 1-day-old and ▲ = 28-day-old). Blue symbols represent foals that remained healthy after intrabronchial infection, gray symbol the foal that developed subclinical pneumonia, and red symbols foals that had clinical pneumonia.

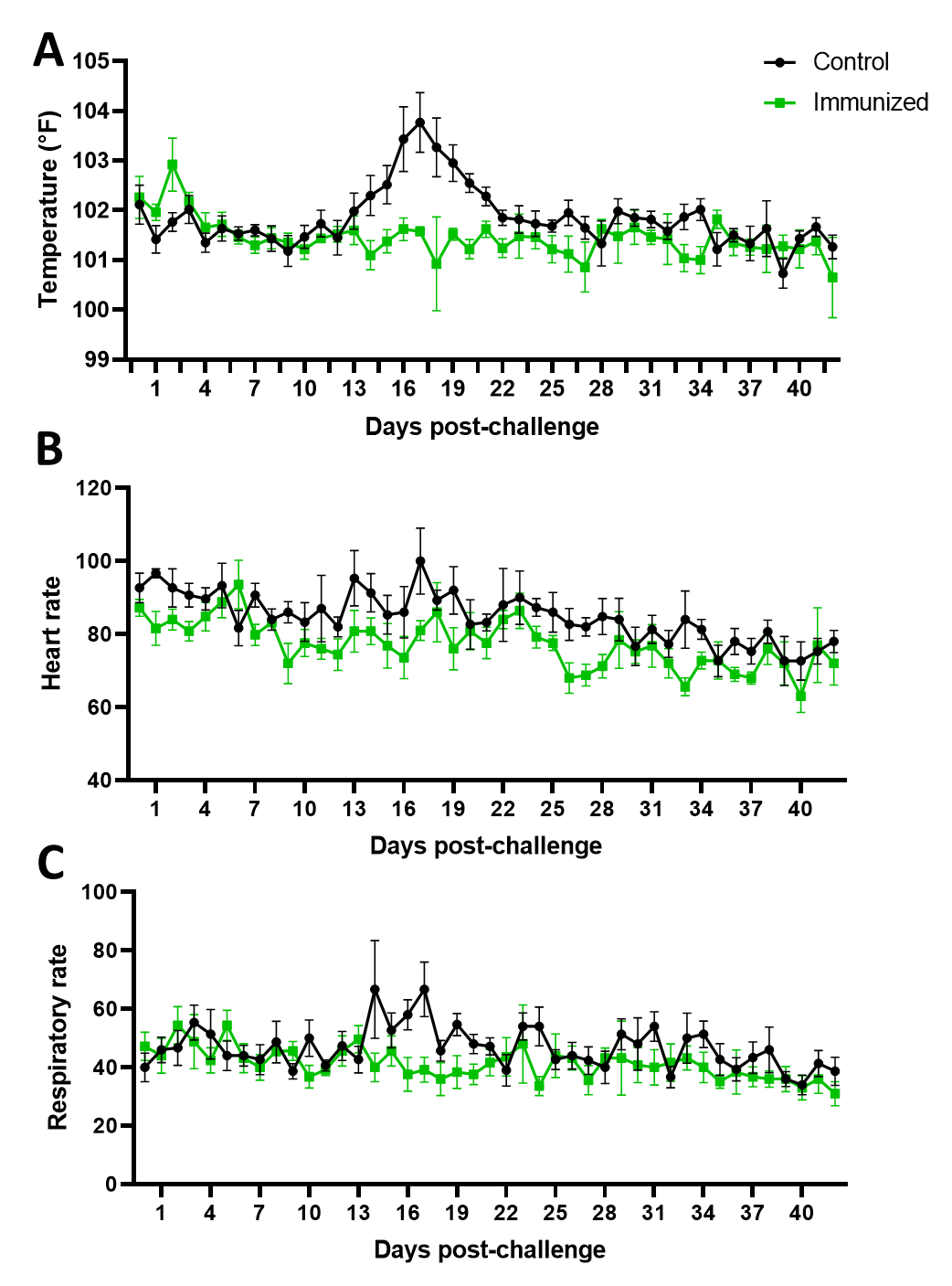

**Fig. S7:** **Clinical monitoring data of foals for 6 weeks following experimental challenge.** Foals were monitored twice daily and vitals recorded. Shown are mean values per group per day, for body temperature in degree Fahrenheit (A), heart rate per minute (B), and respiratory rate per minute (C) data from foals post-challenge. Control foals are shown in black and immunized foals are shown in green. Error bars represent standard error of the mean.
